# Sustained ROS Scavenging and Pericellular Oxygenation by Lignin Composites Rescue HIF-1α and VEGF Levels to Improve Diabetic Wound Neovascularization and Healing

**DOI:** 10.1101/2022.06.18.496670

**Authors:** Jangwook P. Jung, Oluyinka O. Olutoye, Tanuj J. Prajati, Olivia S. Jung, Lane D. Yutzy, Kenny L. Nguyen, Stephen W. Wheat, JoAnne Huang, Benjamin W. Padon, Fayiz Faruk, Sonya S. Keswani, Phillip Kogan, Aditya Kaul, Ling Yu, Hui Li, Shiyanth Thevasagayampillai, Mary E. Guerra, Walker D. Short, Preethi H. Gunaratne, Swathi Balaji

**Author notes:** Corresponding authors: Swathi Balaji, Ph.D. Texas Children’s Hospital and Baylor College of Medicine Feigin Center, C.450.05 1102 Bates Ave. Houston, TX 77030 Jangwook P. Jung, Ph.D. Louisiana State University Department of Biological Engineering 167 E.B. Doran Hall Baton Rouge, LA 70803.

## Abstract

Although delayed wound healing is an important clinical complication in diabetic patients, few targeted treatments are available, and it remains a challenge to promote diabetic wound healing. Impaired neovascularization is one of the prime characteristics of the diabetic phenotype of delayed wound healing. Additionally, increased levels of reactive oxygen species (ROS) and chronic low-grade inflammation and hypoxia are associated with diabetes, which disrupts mechanisms of wound healing. We developed lignosulfonate composites with several wound healing properties, including sustained oxygen release through calcium peroxide nanoparticles and reactive oxygen species (ROS) and free radical scavenging by thiolated lignosulfonate nanoparticles. Sustained release of oxygen and ROS-scavenging by these composites promoted endothelial cell branching and characteristic capillary-like network formation under high glucose conditions *in vitro*. Gene co-expression network analysis of RNA-sequencing results from ECs cultured on lignin composites showed regulation of inflammatory pathways, alongside the regulation of angiogenic hypoxia-inducible factor-1α (HIF-1a) and vascular endothelial growth factor (VEGF) pathways. In vivo, lignosulfonate composite treatment promoted angiogenic growth factor expression and angiogenesis in full thickness skin wounds in diabetic (db/db) mice, a model of delayed wound healing. Treatment of diabetic wounds with lignosulfonate composites also promoted faster epithelial gap closure and increased granulation tissue deposition by day 7 post-wounding, with a higher presence of pro-healing type macrophages. These effects significantly improved tissue repair outcomes by day 14. Our findings demonstrate that lignosulfonate composites promote diabetic wound healing without requiring additional drugs. This highlights the potential of functionalized lignosulfonate for wound healing applications that requires balanced antioxidation and controlled oxygen release.

**Statement of Significance:** The lignosulfonate composites developed in this study offer a promising solution for delayed wound healing in diabetic patients. By effectively addressing key factors contributing to the multifaceted pathophysiology of the diabetic wounds, including impaired neovascularization, increased ROS levels, and chronic inflammation and wound proteolysis, these composites demonstrate significant potential for promoting wound repair and reducing the complications associated with diabetic wounds. The unique combination of pro-angiogenic, oxygen-releasing, ECM remodeling and antioxidant properties in these lignosulfonate-based materials highlights their potential as a valuable therapeutic option, providing a novel approach to diabetic wound healing without the need for additional drugs.

## 1. Introduction

Over eight million people suffer from non-healing wounds every year in the United States [1, 2]. Recurrence of wounds in diabetic patients due to poor healing significantly increases their morbidity, risk for amputations and mortality, and diabetes-related lower extremity complications are among the top 10 leading causes of the global burden of disability [3]. Diabetic wounds have a reduced ability to mount the effective immune response required for pathogen control due to reduced migration of leukocytes, neutrophils, and macrophages to the wound [4, 5]. The accumulation of advanced glycation end products in diabetic wounds increases reactive oxygen species (ROS) formation and reduction of macrophage efferocytosis, thereby impinging their ability to transition to alternatively activated (M2) phenotype and impairing inflammation resolution [6]. This results in increased proteolytic activity along with a decrease in proteolysis inhibitors [7], culminating in insufficient accumulation of granulation tissue and neovascularization. Recent evidence further links excessive production of ROS and/or impaired detoxification of ROS to the pathogenesis of chronic wounds [8–10].

ROS are essential regulators of the wound healing process, facilitaing defense against invading pathogens and, at low levels, being necessary for cellular functions like proliferation, migration, and apoptosis [11]. Thus, ROS production and scavenging is well balanced in normal wound healing Excess ROS are scavenged by enzymes such as superoxide dismutase and antioxidants that regulate the redox environment in healing skin wounds. However, excess ROS accumulation in diabetic wounds disrupts cellular homeostasis, causing non-specific damage to critical cellular components and leading to impairment such as abberant fibroblast collagen synthesis [12] and cell apoptosis. Antioxidants scavenge or neutralize free radical formation, inhibiting the deleterious downstream effects of ROS. This makes them potential therapeutic interventions to combat oxidative stress [13–20]. Strategies including nanoparticles made up of inorganic materials like mesoporous silica, cerium oxide, and fullerene which exhibit antioxidant activities, have been evaluated *in vitro* and in animal models to determine their ability to scavenge free radicals and decrease ROS concentrations, and protect cells against oxidative stress [21–24]. Nevertheless, most antioxidants, taken orally, have limited absorption profile, leading to low bioavailability and insufficient concentrations at the target wound site [25, 26].

Toward the goal of tissue regeneration, ROS-scavenging biomaterials have been identified as promising therapeutic avenues to alleviate oxidative stress in tissue microenvironments [27]. We and others have demonstrated that hydrogels create a provisional wound matrix with good biocompatibility, nutrient supply and swelling that allows the absorption of excess exudates and the maintenance of optimal moist environment [28–32]. These properties facilitate resistance to microbial infections limiting the self-perpetuating cycle of infection-induced oxidative stress in the wounds, and promote antioxidation and ROS scavenging, jump-starting healing by redirecting the wounds from the state of inflammation and ECM proteolysis to the next stages. Furthermore, when considering large wounds, the lack of nutrient supply and oxygen generation beyond the diffusion limit in tissue (100 to 200 μm) has been a major limiting factor for biomaterial-based therapies [33]. In normal wound helaing, hypoxia caused by low oxygen tension in the newly formed wound triggers hypoxia inducible factor 1-α (HIF-1α) mediated adaptive cellular reponses in endothelial cells and fibroblasts to promote angiogenesis and helaing, which is impaired in diabetic donditions. Wound hypoxia significantly hinders healing by disrupting protein synthesis, reducing nutrient supply, weakening immune function and promoting bacterial growth. These deficiencies, coupled with the simultaneous disruption of several pathways involved in the diabetic wound healing response, may in part explain why some current therapies for treating diabetic wounds are not entirely successful. Therefore, we aimed to address this gap by modifying the diabetic wound microenvironment with novel lignin-based composites.

Lignin is a polyphenolic polymer that can isolate pathogens to the site of infection while providing imperviousness to cell walls in vascular plants [34]. The use of lignin in wound-healing dressings is an emerging field [35], with proven effects as an antiviral [36, 37] and antioxidant/immunostimulant [38] . Lignin’s protein adsorption, wound compatibility [39], anti-inflammatory effects via reducing iNOS and IL-1β expression [40], and biocompatibility without toxicity [41] make it promising for wound healing applications.

Not surprisingly, most lignin-based biomaterial applications leverage its antioxidation property [14, 42–50]. To address diabetic wounds, we developed a novel, dual-acting lignin composite with thiolated lignosulfonate (TLS) nanoparticles for antioxidation [51] and lignosulfonate (LS)-based, calcium peroxide nanoparticles (CPO) for controlled oxygen release [52]. This composite not only scavenges ROS but also produces oxygen without harmful byproducts (i.e. H_2_O_2_), creating a corrective, provisional matrix for diabetic wounds. We hypothesized that this lignin composite, with its optimal constituents, would enhance the delivery of beneficial oxygen and antioxidation to infiltrating cells, promoting improved neovascularization and granulation tissue, essential for enhanced healing and tissue regeneration in murine diabetic wounds.

## 2. Materials and Methods

### 2.1. Formation of lignin composites

Our study employed precursors for lignin composites based on our previously published formulations [52]. Briefly, TLS nanoparticles (3 mg/mL) with either CPO (LS nanoparticles formed with CaO_2_ that release O_2_; 4 mg/mL) or CPOc (LS particles formed without CaO_2_ and do not release O_2_ – i.e. CPO’s control; 4 mg/mL) were incorporated in gelatin methacryloyl (GelMA50; mg/mL) with LAP (1 mg/mL; Lithium Phenyl (2,4,6-trimethylbenzoyl) phosphinate). Precursors were cast into custom polydimethylsiloxane (PDMS) wells (**Figure S1**) or injected in db/db mouse wounds followed by UV (ultraviolet) illumination for 60 s (365 nm) to form lignin composites *in vitro* and *in vivo* respectively.

### 2.2. Assessment of capillary-like network formation of MVECs

Mouse dermal microvascular endothelial cells (MVECs) were purchased from Cell Biologics (Boston Bioproducts, Milford, MA). Control MVECs were cultured in Endothelial Cell Medium (Sciencell, Carlsbad, CA) with 5% FBS (fetal bovine serum), while MVECs in the test group were treated with 30 mM dextrose in Endothelial Cell Medium with 5% FBS. MVECs were cultured in 30 mM dextrose for 6 days prior to the tube formation assay and the medium was switched with 1% FBS for cell starvation on the night prior to the assay. On the day of the assay, 50 µL of lignin composite precursors was cast into a custom PDMS well to prevent meniscus formation and facilitate best imaging results. Four groups of lignin composites shown in **Table 1** were cast and then photopolymerized under UV light for 60 s.

**Table 1.** Abbreviation of lignin composites

| Abbreviation | Matrix | Nanoparticles (NPs) |
| --- | --- | --- |
| **UNTX**<sup>1</sup> | None | None |
| **GelMA**<sup>2</sup> | GelMA (50 mg/ml) | None |
| **TLS** | GelMA (50 mg/ml) | TLS (3 mg/ml) |
| **CPOc** | GelMA (50 mg/ml) | TLS (3 mg/ml) and LS-PLGA<sup>3</sup> w/o CaO<sub>2</sub>; 4 mg/ml) |
| **CPO** | GelMA (50 mg/ml) | TLS (3 mg/ml) and LS-PLGA/CaO<sub>2</sub> (4 mg/ml) |

<sup>1</sup>Untreated control for *in vivo* experiments
<sup>2</sup>Untreated control for *in vitro* experiments
<sup>3</sup>PLGA: poly(lactic-*co*-glycolic) acid

MVECs were seeded onto the lignin composites at 5.0 × 10^4^ cells per well and placed in a CO_2_ incubator (37°C/5% CO_2_). A set of randomly selected five images were taken using the inverted fluorescence microscope (Nikon Eclipse Ti2) and NIS Elements Advanced Research Microscope Imaging Software (NIS Elements AR, Nikon) from each lignin composite at every 24 h over 96 h. Angiogenesis Analyzer [53] was used to assess the capillary-like network formation with the five relevant-features of capillary network formation and angiogenesis quantified.

### 2.3. Assessment of HIF-1α and VEGF in culture medium of MVECs

Following the network formation of MVECs, supernatant was collected at every 24 h over 96 h, and protease and phosphatase inhibitors were added to the medium before freezing at -80°C. Expression levels of murine vascular endothelial growth factor (VEGF) (Quantikine ELISA (enzyme-linked immunosorbent assay) kit, R&D Systems, Minneapolis, MN, USA) and HIF-1α (SimpleStep ELISA kit, Abcam ab275103, Cambridge MA, USA) in the culture medium were determined using ELISA.

### 2.4. RNA sequencing (RNA-seq) of MVECs and analysis

RNA was extracted from MVECs seeded on the lignin composites using Trizol, and the aqueous phase was subsequently purified using an RNA mini kit (PureLink RNA Mini kit). Bulk RNA sequencing was conducted at the University of Houston Sequencing Core. Further details are in Supplementary Methods. The variability in the data was identified using Principal Component Analysis (PCA). R package WGCNA (Weighted Gene Co-Expression Network Analysis) was utilized to create the gene co-expression network [54]. The network was assembled using a soft thresholding power (β) of 0.9. Modules with high degrees of similarity were merged into a single module at a threshold of 0.4. To determine which module had the strongest connection to the samples, a Pearson correlation analysis was performed to investigate the relationships between gene modules. Candidate genes obtained from WGCNA were submitted to STRING and ClueGO plugin to Cytoscape [55] for visualization.

### 2.5. In vivo model of diabetic wound healing

The db/db mouse model of diabetic (type II) wound healing was used to determine the effect of lignin composites on angiogenesis and wound healing *in vivo*. This murine model is characterized by a delayed wound healing phenotype with reduced neovascularization [56] and increase in ROS and inflammation. All procedures were approved by the Institutional Animal Care and Use Committee. Eight-week-old female, diabetic B6.BKS(D)-*Lepr^db^*/J (db/db) mice were obtained from Jackson Laboratories (Bar Harbor, ME). Two full thickness 6 mm circular excisional splinted wounds were created on the bilateral dorsal flanks of each animal, leaving the underlying muscle (panniculus carnosus) intact, as previously described [56]. Treatment groups included UNTX, or wounds treated with lignin composites TLS, CPOc and CPO (**Table 1**) which were injected into the wounds as a one-time dose (50 µl) immediately after wounding.

Animals were euthanized at days 7 and 14, and wound tissues were harvested, and fixed and paraffin embedded or flash frozen and stored at -80°C for further analysis.

### 2.6. Immunohistochemistry

Immunohistochemistry staining was performed on wound sections using Dako Auto-stainer Link 48 (DakoLink version 4.1, edition 3.1.0.987; Agilent, Santa Clara, CA) as described previously [52]. Primary antibodies against CD31 for endothelial cells (ab28364; 1:100; Abcam, Cambridge, MA), VEGF (MA5-3208; 1:300, Thermofisher Scientific, Waltham, MA) and HIF-1a (ab179483; 1:100; Abcam, Cambridge, MA), CD45 for pan leukocytes (ab10558; 1:5000; Abcam, Cambridge, MA), F4/80 for pan-macrophages (ab111101; 1:100: Abcam, Cambridge, MA) were detected by EnVision+System-HRP (3,3 Diaminobenzidine, DAB) kits (Dako North America, Carpinteria, CA) and hematoxylin counter staining. Histology slides were imaged with Leica DM 2000® with Leica Application Suite X® version 3.0.4.16529. Immunofluorescence staining was performed using primary antibodies against CD206 (ab64693, 1:500, Abcam, Cambridge, MA) and Arginase-1 (PA529645, 1:500, Thermo Fisher, Waltham, MA) followed by Alexafluor^TM^ (ThermoFisher, Waltham, MA) secondary antibodies to co-label macrophages.

### 2.7. Quantification of HIF-1α and VEGF in the wound homogenates

To determine protein expression of VEGF and HIF-1α in the db/db mice wounds treated with lignin composites, wounds from day 7 post wounding were harvested, and homogenized using RIPA buffer (ThermoFisher Scientific 89900, Waltham, MA, USA). The digested samples were centrifuged, and the supernatant was used in the quantification of expression levels of murine VEGF and HIF-1ausing Quantikine ELISA as described above.

### 2.8. Statistical analyses

Statistical comparisons between groups were performed by one-way ANOVA (analysis of variance). For multiple comparisons, one-way ANOVA with Tukey’s or Dunnett’s *post hoc* comparison was carried out. p values <0.05 were considered statistically significant. All bar graphs represent mean ± standard deviation (SD).

## 3. Results

### 3.1. Endothelial capillary-like network formation improved on dual-acting lignin composites

Following 6 days of treatment with 30 mM glucose in 2D cultures, MVECs were seeded onto four different types of lignin composites, GelMA, TLS, CPOc and CPO (see **Table 1** for details of abbreviations) up to 96 h. Phase contrast micrographs were taken and analyzed using Angiogenesis Analyzer [53]. As shown in **Figure 1**, lignin composites supported the growth and capillary-like network formation of MVECs. Five features were selected from the Angiogenesis Analyzer including number of extremities, number of nodes, number of junctions, number of branches and average mesh sizes (**Figure 2**). Lignin composites improved the characteristic capillary-like angiogenesis features under both normal and oxidative stress conditions. Number of extremities, number of branches, and mean mesh size are significantly higher in CPO lignin composite (high glucose) in comparison to CPOc. All lignin composites exhibited higher numbers of nodes and junctions regardless of the presence of the high glucose condition.

**Figure 1.**
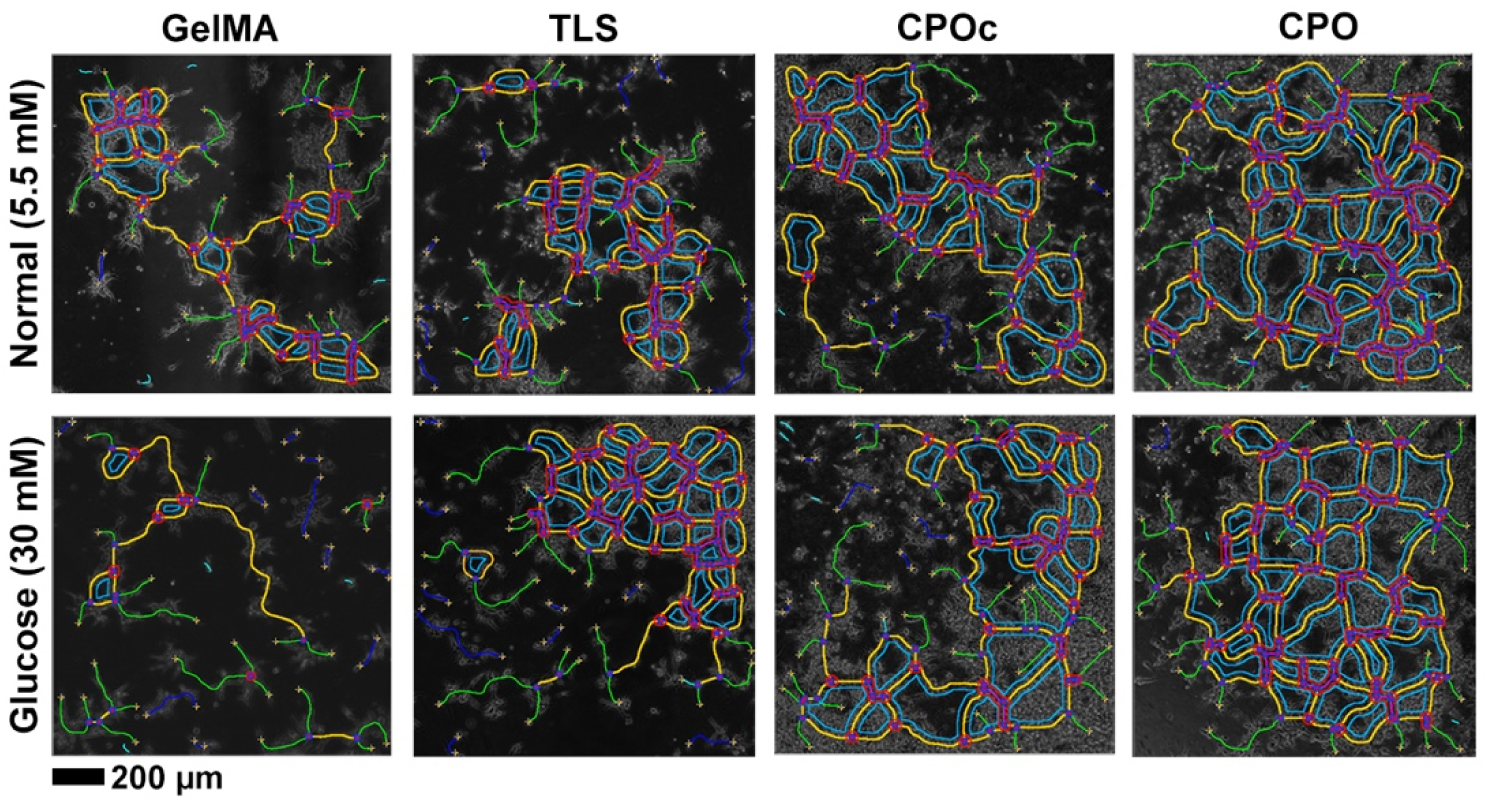
Results from Angiogenesis Analyzer [53] for the micrographs taken after 96 h of MVEC culture initiation on the different lignin composites and controls, GelMA, TLS, CPOc and CPO (see **Table 1** for details of compositions) Green - branches; Cyan - twigs and small isolated segments; Magenta - segment; Yellow/Orange - master segments; Sky blue - mesh; Dark blue - isolated elements.

**Figure 2.**
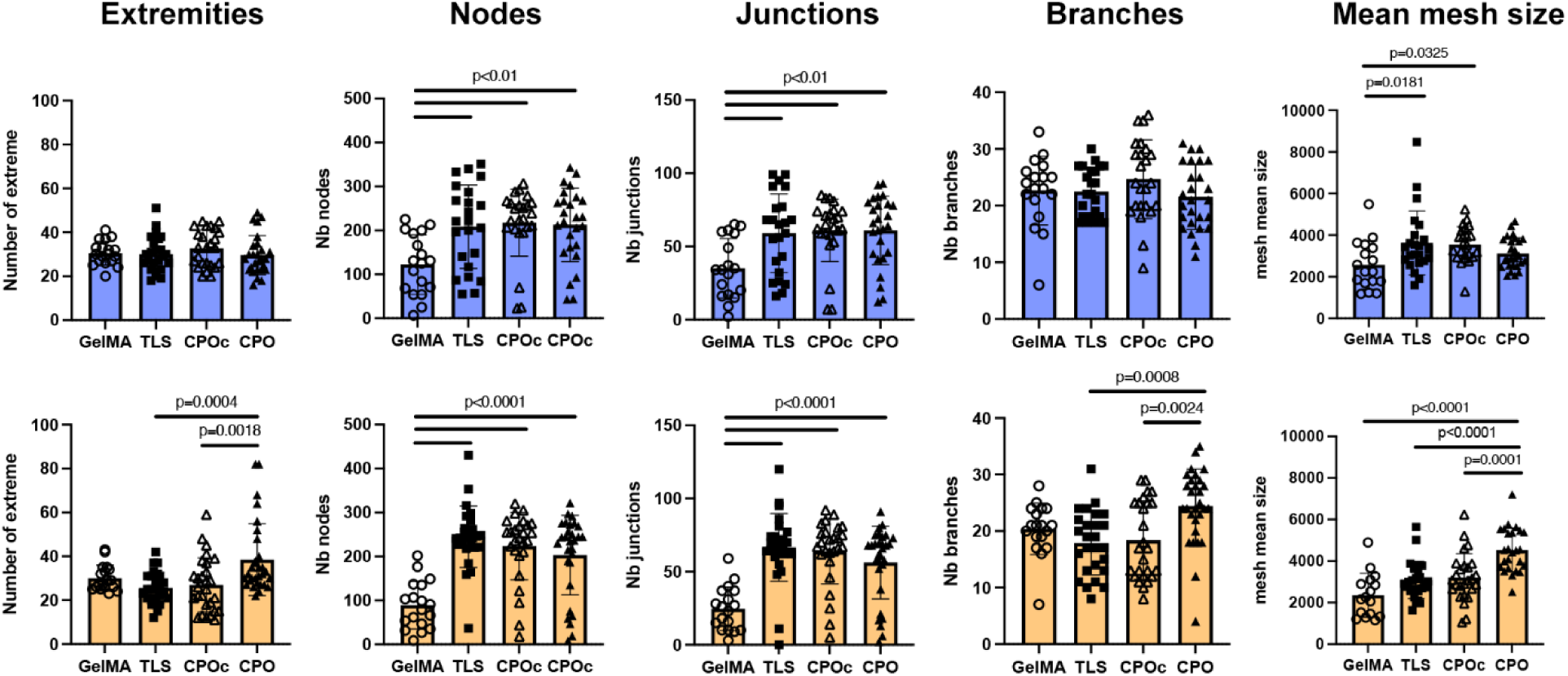
Five angiogenic features are analyzed to assess the effect of lignin composites on MVECs in two different conditions (blue, normal glucose 5.5 mM; orange, high glucose 30 mM): number of extremities, number of nodes, number of junctions, number of branches and average mesh sizes are analyzed. Mean±SD, n=6-9 independent samples, 3 images/composite were analyzed. p values by one way ANOVA and Tukey’s *post hoc* tests.

### 3.2. Gene co-expression network analysis identified the co-expression of Hif1α and Vegf in MVECs cultured on lignin composites

Bulk RNA-seq of samples revealed distinct principal components (PCs) in **Figure 3A**. PC1 distinguished lignin composites from GelMA, while PC2 further separated GelMA samples cultured with and without high glucose. To visualize gene co-expression network, we applied our samples to WGCNA (other relevant modules are in **Figure S2** with all 12 samples subject to WGCNA). Of 29 modules identified, the turquoise module containing 5,073 genes was selected for distinguished module-sample relationship (**Figure 3B**). As demonstrated in **Figure 3C**, CPO composites facilitated the expression of *Hif1α* in the pool of top 25 hub genes. In enrichment analysis, *Vegf* associated pathways were significantly upregulated (**Figure 3D** and **3E**) as a part of “cytokine production involved in inflammatory response” network.

**Figure 3.**
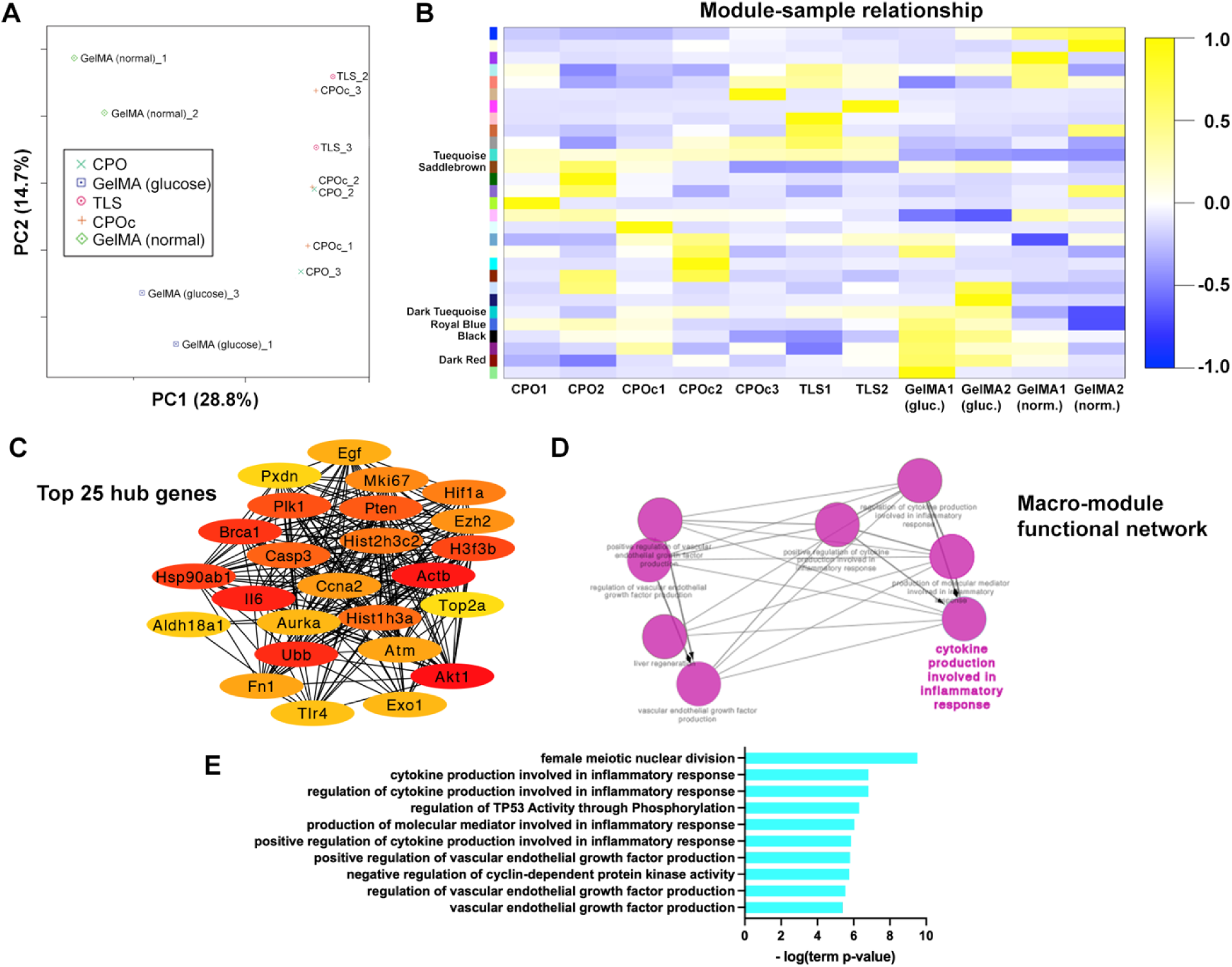
RNA-seq analysis of MVECs cultured on lignin composite TLS, CPO or CPOc along with GelMA controls (with and without high glucose medium conditions). All samples were subject to (A) PCA and (B) WGCNA module-sample relationship. In enrichment analysis, the turquoise module (containing 5,073 genes) included hub genes (C) relevant to *Hif1*α, including the most significant functional categories including *Vegf* (D) and (E) identified by ClueGO Cytoscape plugin.

### 3.3. VEGF and HIF-1α expression of MVECs was altered in vitro in lignin composites

VEGF and HIF-1α, two important growth factors involved in wound neovascularization, are highly susceptible to ROS-mediated variations in the wound milieu. These variations ultimately guide angiogenic responses. Hence, we sought to determine their expression in MVECs cultured on lignin composites under normal and high glucose conditions. At 96 h, HIF-1a was expressed highest in the media of high glucose MVECs cultured on the CPO lignin composites (**Figure 4A**). HIF-1α responses on other lignin composites were comparable, except for GelMA, which is likely due to the very small variance within groups. Unlike HIF-1α, differences in VEGF expression became apparent from 24 h. MVECs cultured under high glucose conditions showed a significantly higher VEGF expression compared to normal glucose media conditions on GelMA controls. Under normal glucose media conditions, VEGF expression of MVECs cultured on lignin composites was not different from GelMA controls. Notably, VEGF expression of MVECs under high glucose media conditions when cultured on lignin composites with antioxidation (TLS) began to show attenuation to normal expression levels (**Figure 4B**), with CPO lignin composites with dual functionality of antioxidation and oxygenation showing the most attenuation of VEGF levels.

**Figure 4.**
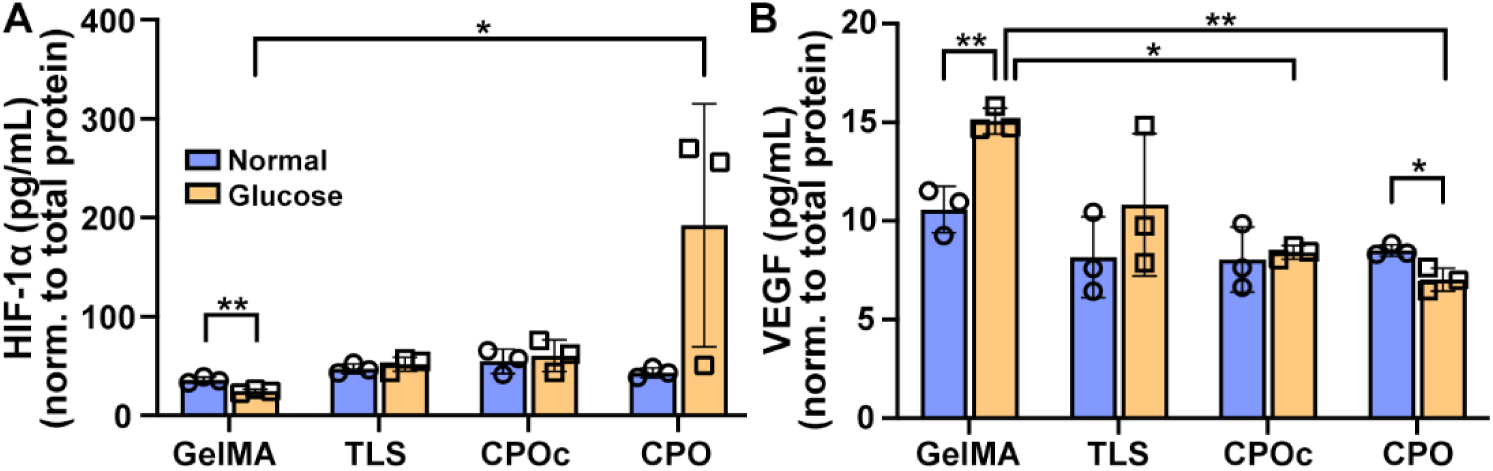
Analysis of levels of HIF-1α at 96 h (A) and VEGF at 24 h (B) collected in MVEC culture medium containing either normal (5.5 mM) glucose or high (30 mM) glucose. Mean±SD, n=3. *p<0.05 and **p<0.01 considered significant by one way ANOVA Tukey’s *post hoc* tests.

### 3.4. Antioxidant, oxygen-generating dual-acting lignin composites improved granulation tissue and capillary lumen formation in db/db wounds at day 7

We evaluated wound healing in 8-week-old db/db mice with blood glucose >3.5 mg/mL with 6 mm wounds (**Figure 5A**) treated with lignin composites TLS, CPO or CPOc immediately after wounding and compared to healing in untreated wounds (UNTX). Hematoxylin and Eosin (H&E) staining of wound sections at day 7 showed no excessive contraction of the wounds with the lignin composites or any other inadvertent events **(Figure 5B)**. Re-epithelialization was noted in all wounds. To accurately assess the epithelial gap amongst all treatment conditions, cK14 (cytokeratin 14) staining of wound sections was performed to stain the keratinocytes (**Figure S3**), which showed least gap between the encroaching margins or open wound in CPO treated wounds compared to other treatments (UNTX: 4.79 ± 1.37 mm, TLS: 4.54 ± 1.35 mm, CPOc: 3.6 ± 1.15 mm, CPO: 3.32 ± 0.60 mm, p=0.054 UNTX vs. CPO) (**Figure 5C**). In contrast, wounds treated with CPO appeared to have more granulation tissue present at day 7 (UNTX: 0.5 ± 0.1 mm^2^, TLS: 0.6 ± 0.4 mm^2^, CPOc: 0.4 ± 0.2 mm^2^, CPO: 1.7 ± 0.8 mm^2^) when compared to other wounds (**Figure 5D**). These data indicate a notable trend with improvement in wound morphology in dual-acting, CPO lignin composite treated wounds.

**Figure 5.**
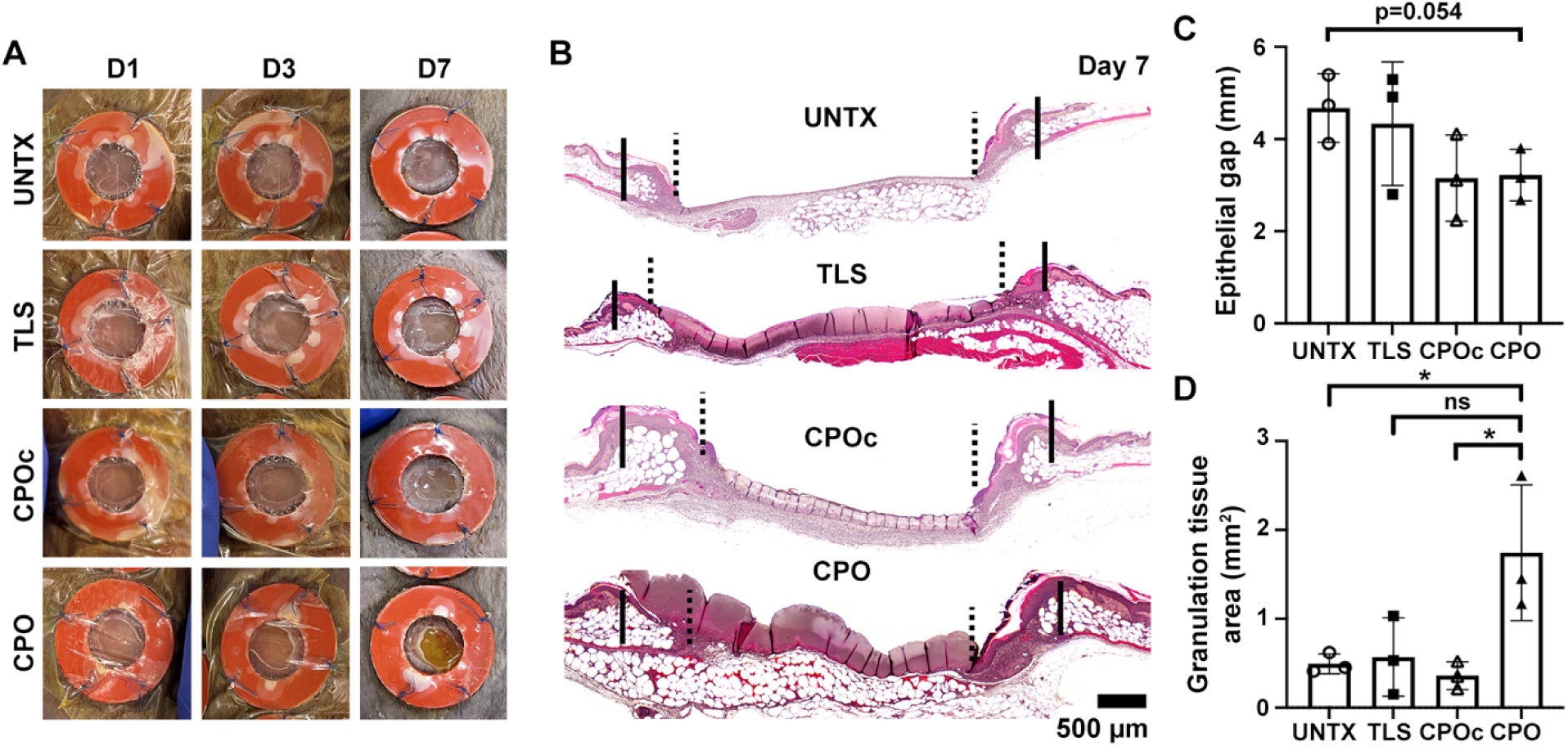
(A) Gross images taken at indicated intervals post-wounding show healing progression of db/db mouse wounds treated with lignin composites TLS, CPO or CPOc compared to UNTX wounds. (B) Representative H&E staining of wounds at post-wounding day 7, Scale bar = 500 µm. Quantification of epithelial gap (C) and granulation tissue area (D) at day 7 post wounding. Mean±SD, n=3 wounds per treatment group, *p<0.05 by one way ANOVA Tukey’s *post hoc* tests.

CD31 staining of wound sections at day 7 showed a significant increase in capillary lumen density in CPO treated wounds (11 ± 1.2 vessels/HPF), when compared to UNTX (6 ± 0.1 vessels/HPF, p<0.01), TLS (6 ± 1.7 vessels/HPF, p=0.02), and CPOc (6 ± 2.4 vessels/HPF, p=0.049) **(Figure 6A and 6D)**.

**Figure 6.**
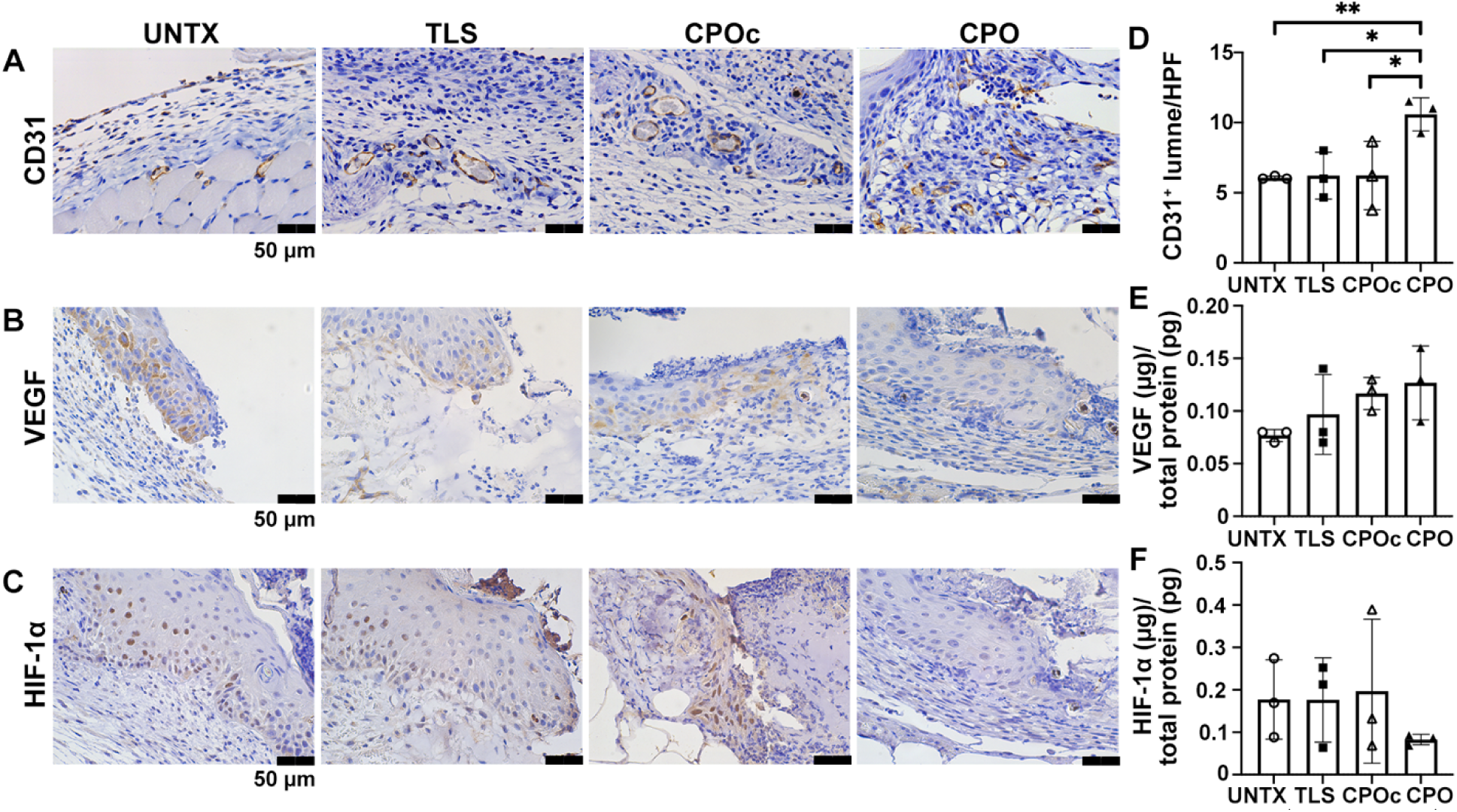
Representative panels from db/db mouse wounds treated with lignin composite TLS, CPO or CPOc compared to UNTX wounds. Staining of day 7 wound sections with antibodies against CD31 (A), VEGF (B) and HIF-1a (C), Scale bar = 50 µm. (D) CD31 staining of wound sections showed a significant increase in lumen density per HPF in CPO wounds at day 7 group. Day 7 wound ELISA revealed an increasing trend in VEGF expression (E) and decreased HIF-1a expression (F). Mean±SD, n=3 wounds per treatment, *p<0.05 and **p<0.01 by one way ANOVA Tukey’s *post hoc* tests.

Interestingly, we noted striking differences in VEGF and HIF-1α expression with immunohistochemistry at the leading epidermal margins in CPO vs. other treatments. Consistent with our previous work on normal wound healing in murine skin wounds [52], db/db mice leading epidermal edges in UNTX wounds displayed abundant VEGF and HIF-1α expression at day 7 post wounding, which is upregulated in response to injury-induced oxidative stress **(Figure 6B and 6C)**. Similar expression patterns were noted in TLS and CPOc wounds. Strikingly, the oxygen generating lignin composite CPO did not elicit this response at day 7. While immunohistochemical staining of VEGF was reduced in the epidermis, quantification of VEGF expression in the homogenized wound granulation tissue using ELISA showed an increase in the VEGF expression at day 7 in the wound bed **(Figure 6E)**, suggesting dermal angiogenesis is promoted by CPO lignin composites. HIF-1a quantification of the homogenized wounds showed reduced expression in CPO wounds at day 7 as compared to UNTX, TLS and CPOc wounds **(Figure 6F)**.

In addition, we did not see any significant difference in the expression of CD45^+^ (pan-leukocyte) cells across all tested lignin composites when compared to untreated wounds at day 7. CD45 staining displayed a slight decrease in CPO-treated wounds (13 ± 10 % CD45^+^ cells/HPF) compared to untreated (24 ± 10 % CD45^+^ cells/HPF, p=0.26) and CPOc (16 ± 9 %CD45^+^ cells/HPF, p=0.72) conditions, but that is not the case for TLS (10 ± 2 % CD45^+^ cells/HPF, p=0.60) **(Figure 7A and 7D)**. F4/80 staining displayed a significant decrease of macrophages in CPO-treated wounds compared to TLS-treated wounds (17 ± 2 % F4/80^+^cells/HPF vs. 41 ± 6 % F4/80^+^cells/HPF, p<0.01). F4/80 staining was decreased in CPO-treated wounds compared to both untreated (26 ± 12 % F4/80^+^cells/HPF, p=0.26) and CPOc-treated (20 ± 10 % F4/80^+^cells/HPF) wounds, however these differences did not reach significance **(Figure 7B and 7E)**, indicating minimal inflammatory response from CPO or CPOc nanoparticles in lignin composites. In line with our hypothesis, we quantified the expression of CD206 and Arg1 in macrophages. As demonstrated in **Figure 7C and 7F**, the percentage of CD206^+^ Arg1^+^ of the total CD206^+^ macrophages progressively increased with antioxidation and oxygenation, with the highest expression observed in CPO-treated wounds in comparison to UNTX (74 ± 6.3 % CD206^+^ Arg1^+^ of CD206^+^ vs. 47 ± 12 %, p<0.05).

**Figure 7.**
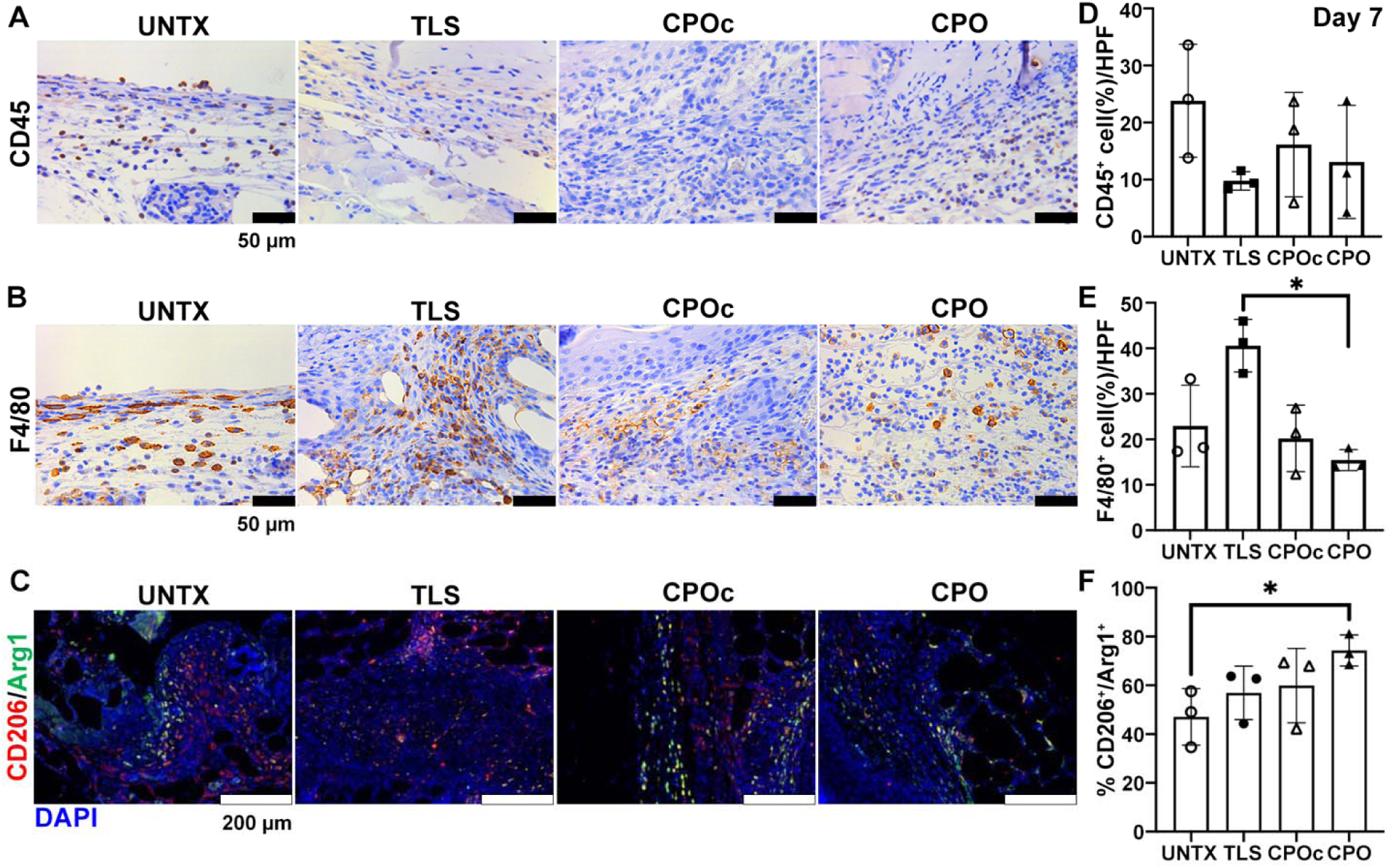
Inflammatory panel in db/db mouse wounds treated with lignin composite TLS, CPO or CPOc compared to control UNTX. Representative images of stained day 7 wound sections with antibodies against CD45 (A), F4/80 (B) and CD206/Arginase1 (C). Scale bar = 50 µm for (A) and (B); scale bar = 200 µm for (C). Quantification of % of CD45^+^ (C) and F4/80^+^ (D) cells in HPF in the wound sections showed no significant increase in inflammatory markers in CPOc or CPO wounds at day 7. Mean±SD, n=3 wounds per treatment, *p<0.05 by one way ANOVA Tukey’s *post hoc* tests. Quantification of CD206^+^/Arg1^+^ macrophages among the CD206^+^ cells. Mean±SD, n=3 wounds per treatment. *p<0.05 by one way ANOVA with Dunnett’s multiple comparison tests.

To determine the effect of the lignin composite treatment on remodeling in db/db wounds, we followed the wounds to later time points, and assessed the effect of treatments at 14 days post wounding. At day 14, we noted visibly improved healing in CPOc and CPO wounds from gross images, which was supported by the presence of a robust granulating wound bed in representative hematoxylin and eosin-stained wounds sections **(Figure S4A).** Day 14 wounds treated with CPO (24 ± 4 CD31^+^ lumens/HPF) also displayed increased lumen density with CD31 staining compared to TLS (12 ± 5 CD31^+^ lumens/HPF) and CPOc (14 ± 2 CD31^+^ lumens/HPF) but did not approach significance compared to UNTX (19 ± 0.33 CD31^+^ lumens/HPF) **(Figure S4B** and **S4C)**.

## 4. Discussion

We sought to determine whether lignin-based composites promote diabetic wound healing by mitigating the effects of ROS. We tested whether antioxidant and oxygen-generating lignin composites can enhance endothelial cell functions under hyperglycemic conditions. We noted a significant improvement in MVEC network formation with CPO **(Figure 1)**, suggesting that oxygen-generating lignin composites might correct cellular dysfunctions in diabetic ECs. However, recent reports suggest that an *in vitro* acute hyperglycemic model may not be ideal for mechanistic studies of the effects of diabetes on endothelial cell responses due to a phenomenon called “hyperglycemic memory” [57–61]. Experimental evidence suggests that oxidative stress results in sustained activation of antiangiogenic, proinflammatory pathways in MVECs even after glycemia is normalized (i.e., the phenotype of endothelial cells becomes altered permanently, although the exact mechanisms are not well understood), as shown in our previous studies [31]. In addition, previous studies [62, 63] have shown an increase in VEGF expression in diabetic endothelial cells and endothelial cells under excessive ROS, which was also noted in our cell culture experiments with MVEC under high concentration of glucose. Culturing MVECs on CPO lignin composites reversed these effects induced by high concentration of glucose [62]. These findings suggest a need for novel therapies that can reverse endothelial cell hyperglycemic memory and stimulate neovascularization. Our lignin composite design could fill this gap.

Interestingly, differential gene expression (DGE) and subsequent gene ontology (GO) analysis revealed that all lignin-containing samples (e.g., CPO, CPOc, and TLS) exhibited regulation of membrane potential (GO:0042391) (**Figure S5**). In contrast, GelMA controls, both with and without glucose, did not show this pattern (**Figure S5G**). These findings suggest that MVECs cultured on lignin composites actively adapt to oxidative stress by modulating membrane potential. **Figure S5C** indicates no DGE differences between CPOc and TLS, supporting our design criterion that CPOc is practically identical to TLS without oxygenation. Due to relatively low numbers of significant DGEs, further optimization and increased sample sizes are warranted for future investigations. Conversely, **Figures S5D-F** demonstrate that each individual component of lignin composites, unlike GelMA under oxidative stress (high glucose, 30 mM), positively contributes to membrane potential regulation. Moreover, the presence of oxidative stress impacted DGE (**Figure S5G**).

Motivated to better understand these correlations, we applied WGCNA and identified HIF-1α as a top hub gene. VEGF emerged as a key factor in enriched pathways identified through bulk RNA-seq analysis (Figure 3E). However, the number of samples used for WGCNA analysis (Figures 3 and **S2**) falls short of the recommended minimum of 15. Consequently, only the turquoise module is relevant to both HIF-1α and VEGF. This limitation suggests the necessity for further studies with more samples.

We further validated the expression of these proteins in MVEC cultures on lignin composites using ELISA (Figure 4). Surprisingly, while HIF-1α expression significantly increased over 96 h in MVEC cultures on CPO lignin composites with high glucose medium compared to controls like GelMA and CPOc, VEGF expression did not exhibit a similar increase in the first 24 h. Instead, MVECs secreted more VEGF in the presence of oxidative stress, and this appeared to be mitigated by the presence of functional lignosulfonate nanoparticles alone (TLS) or together (CPO), as shown in Figure 4B. Under oxidative stress, high VEGF levels are often associated with leaky, immature, poorly perfused vessels [64]. The topology of the functional network (Figure 3D) and enrichment analysis (Figure 3E) partially supported this finding, suggesting VEGF, although significant, cannot solely explain the primary inflammatory response. Multiple time points and single-cell RNA-sequencing would provide a more extensive understanding of how CPO lignin composites affect VEGF expression.

Next, we then harvested wounds from 8-week-old db/db mice with 6 mm wounds treated with lignin composites. Diabetic wounds in db/db mice are marked by decreased granulation tissue formation [65]. However, we observed that granulation tissue was significantly improved at day 7 in CPO lignin composite-treated wounds. This robust granulation issue formation in the CPO lignin composites suggests that scavenging free radicals and ROS via lignosulfonate in the wound rescues the alterations caused by hyperglycemic memory and restores robust granulation tissue formation. While we do not know yet fully understand the precise molecular mechanism underlying this process, it is an area of our further research. A deeper understanding of the pathways involved in this restoration of robust granulation tissue may provide insight into how to improve diabetic wounds and decrease the incidence of wound recurrence.

Leading edges of wounds showed HIF-1α induction. Interestingly, in CPO composite treated wounds containing oxygen-generating nanoparticles, HIF-1α increase was not significant at the edges. However, the presence of granulation tissue confirms ongoing wound healing. This suggests that oxygenation of the wounds change hypoxia-induced pathways in diabetic wounds and may reduce HIF-1α expression and its downstream signaling. Further understanding of these pathways leading to HIF-1α expression in diabetic mice treated with lignin composites will be the focus of future projects. VEGF, which is well-known for its proangiogenic abilities [66, 67], also plays a significant role in cutaneous wound healing. Interestingly, despite the known connection between HIF-1α and VEGF [68–70], we see that VEGF expression is increased in the CPO-treated wounds (Figure 6B**/E**), suggesting that CPO scavenges ROS in the wound and may increase VEGF via pathways independent of HIF-1α (Figure 6C**/F)**. The mechanism by which this occurs is not well described and could be a future area of study, although, increased oxygen levels in diabetic wounds have previously been shown to increase wound VEGF levels. For example, in the rat wound model, hyperbaric oxygen treatment resulted in increased VEGF in wound fluid [71]. We have shown that this increase in VEGF in CPO-treated wounds is accompanied by expected increase in capillary lumen density and improved wound healing. Further understanding of the role CPO lignin composites played in VEGF increase will help improve the design of our engineered scaffolds to increase their wound healing efficacy.

Although expression levels of CD45^+^, F4/80^+^, and CD206^+^/Arg1^+^ macrophage markers were relatively similar, we observed some non-significant differences. The decrease in CD45 (associated with leukocytes) and increase in F4/80 (associated with macrophages) suggest a shift from a pro-inflammatory (M1) to an anti-inflammatory (M2) macrophage phenotype. This shift is more pronounced with TLS (Figures 7D and **7E**). The observed changes may indicate that TLS and CPO lignin composites have immunosuppressive effects, reducing immune cell infiltration (CD45) and promoting an anti-inflammatory macrophage phenotype (CD206/Arginase 1, Figure 7F). Incorporating additional controls lacking TLS nanoparticles in each composite formulation, can provide a clearer understanding of TLS’s specific contribution to diabetic wound healing when combined with CPO groups.

A limitation of this study is that we use the db/db mouse as a model for delayed diabetic wound healing. We created excisional wounds manually, not spontaneously as in human diabetic wounds/ulcers. Studies in porcine models with skin architecture similar to that of humans will yield more relevant translational data, which will be a focus of our future studies.

## 5. Conclusions

We have demonstrated that ROS scavenging by the constitutes of lignin composites, coupled with locoregional oxygen generation, decreases HIF-1a expression, increases VEGF expression and increases granulation tissue formation in a wounded db/db mouse model. These data support the well-established notion that oxidative stress causes impaired wound healing, suggesting a potential solution of improving diabetic wound healing by increasing oxygen levels in the wound via free radical scavenging by our dual-acting CPO lignin composite. Future studies will focus on sampling-efficient optimization of the CPO lignin composite to further improve the diabetic wound microenvironment, with the goal of transitioning to large animal studies and ultimately translating to human diabetic wounds.

## CRediT authorship contribution statement

**Jangwook P. Jung:** Conceptualization, Data Curation, Formal Analysis, Funding Acquisition, Investigation, Methodology, Project Administration, Resources, Supervision, Validation, Visualization, Writing – Original Draft Preparation, Writing – Review & Editing. **Oluyinka O. Olutoye II:** Data Curation, Investigation. **Tanuj J. Prajapati:** Data Curation, Investigation, Visualization. **Olivia S. Jung:** Data Curation, Investigation, Methodology. **Lane D. Yutzy:** Investigation, Methodology. **Kenny L. Nguyen:** Investigation, Methodology. **Stephen W. Wheat:** Investigation, Methodology. **JoAnne Huang**: Investigation, Visualization. **Benjamin W. Padon:** Data Curation, Investigation. **Fayiz Faruk:** Data Curation, Investigation. **Sonya S. Keswani:** Investigation, Methodology. **Phillip Kogan:** Investigation, Methodology. **Aditya Kaul:** Investigation, Methodology. **Ling Yu:** Investigation, Methodology. **Hui Li:** Investigation, Methodology. **Mary E. Guerra:** Writing – Review & Editing. **Walker D. Short:** Formal Analysis, Writing – Review & Editing. **Shiyanth Thevasagayampillai** and **Preethi Gunaratne:** RNA-sequencing and data analysis. **Swathi Balaji:** Conceptualization, Data Curation, Formal Analysis, Funding Acquisition, Investigation, Methodology, Project Administration, Resources, Supervision, Validation, Visualization, Writing – Original Draft Preparation, Writing – Review & Editing.

## References

[1] C.K. Sen, Human Wounds and Its Burden: An Updated Compendium of Estimates, Adv Wound Care (New Rochelle) 8(2) (2019) 39–48.

[2] S.R. Nussbaum, M.J. Carter, C.E. Fife, J. DaVanzo, R. Haught, M. Nusgart, D. Cartwright, An Economic Evaluation of the Impact, Cost, and Medicare Policy Implications of Chronic Nonhealing Wounds, Value Health 21(1) (2018) 27–32.

[3] New CDC report: More than 100 million Americans have diabetes or prediabetes, Diabetes growth rate steady, adding to health care burden, Centers for Disease Control and Prevention, 2017.

[4] M. Delamaire, D. Maugendre, M. Moreno, M.C. Le Goff, H. Allannic, B. Genetet, Impaired leucocyte functions in diabetic patients, Diabet Med 14(1) (1997) 29–34.

[5] J. Wysocki, B. Wierusz-Wysocka, A. Wykretowicz, H. Wysocki, The influence of thymus extracts on the chemotaxis of polymorphonuclear neutrophils (PMN) from patients with insulin-dependent diabetes mellitus (IDD), Thymus 20(1) (1992) 63–7.

[6] A. Das, K. Ganesh, S. Khanna, C.K. Sen, S. Roy, Engulfment of apoptotic cells by macrophages: a role of microRNA-21 in the resolution of wound inflammation, J Immunol 192(3) (2014) 1120–9.

[7] N.J. Trengove, M.C. Stacey, S. MacAuley, N. Bennett, J. Gibson, F. Burslem, G. Murphy, G. Schultz, Analysis of the acute and chronic wound environments: the role of proteases and their inhibitors, Wound Repair Regen 7(6) (1999) 442–52.

[8] M. Schafer, S. Werner, Oxidative stress in normal and impaired wound repair, Pharmacol Res 58(2) (2008) 165–71.

[9] C. Dunnill, T. Patton, J. Brennan, J. Barrett, M. Dryden, J. Cooke, D. Leaper, N.T. Georgopoulos, Reactive Oxygen Species (ROS) and Wound Healing: the Functional Role of ROS and Rmerging ROS-Modulating Technologies for Augmentation of the Healing Process, Int Wound J 14(1) (2017) 89–96.

[10] A. Nouvong, A.M. Ambrus, E.R. Zhang, L. Hultman, H.A. Coller, Reactive oxygen species and bacterial biofilms in diabetic wound healing, Physiol Genomics 48(12) (2016) 889–896.

[11] C.K. Sen, S. Roy, Redox signals in wound healing, Biochim Biophys Acta 1780(11) (2008) 1348–61.

[12] D.A. Siwik, P.J. Pagano, W.S. Colucci, Oxidative Stress Regulates Collagen Synthesis and Matrix Metalloproteinase Activity in Cardiac Fibroblasts, Am J Physiol Cell Physiol 280(1) (2001) C53–60.

[13] D. Sacks, B. Baxter, B.C.V. Campbell, J.S. Carpenter, C. Cognard, D. Dippel, M. Eesa, U. Fischer, K. Hausegger, J.A. Hirsch, M. Shazam Hussain, O. Jansen, M.V. Jayaraman, A.A. Khalessi, B.W. Kluck, S. Lavine, P.M. Meyers, S. Ramee, D.A. Rüfenacht, C.M. Schirmer, D. Vorwerk, Multisociety Consensus Quality Improvement Revised Consensus Statement for Endovascular Therapy of Acute Ischemic Stroke, Int J Stroke 13(6) (2018) 612–632.

[14] D. Kai, K. Zhang, L. Jiang, H.Z. Wong, Z. Li, Z. Zhang, X.J. Loh, Sustainable and Antioxidant Lignin–Polyester Copolymers and Nanofibers for Potential Healthcare Applications, ACS Sustainable Chemistry & Engineering 5(7) (2017) 6016–6025.

[15] S. Hajihashemi, S. Rajabpoor, I. Djalovic, Antioxidant potential in Stevia rebaudiana callus in response to polyethylene glycol, paclobutrazol and gibberellin treatments, Physiol Mol Biol Plants 24(2) (2018) 335–341.

[16] R. Javed, M. Ahmed, I.U. Haq, S. Nisa, M. Zia, PVP and PEG doped CuO nanoparticles are more biologically active: Antibacterial, antioxidant, antidiabetic and cytotoxic perspective, Mater Sci Eng C Mater Biol Appl 79 (2017) 108–115.

[17] W. Zeng, J. Xiao, G. Zheng, F. Xing, G.L. Tipoe, X. Wang, C. He, Z.Y. Chen, Y. Liu, Antioxidant Treatment Enhances Human Mesenchymal Stem Cell Anti-Stress Ability and Therapeutic Efficacy in an Acute Liver Failure Model, Sci Rep 5 (2015) 11100.

[18] G. Choe, S.W. Kim, J. Park, J. Park, S. Kim, Y.S. Kim, Y. Ahn, D.W. Jung, D.R. Williams, J.Y. Lee, Anti-Oxidant Activity Reinforced Reduced Graphene Oxide/Alginate Microgels: Mesenchymal Stem Cell Encapsulation And Regeneration of Infarcted Hearts, Biomaterials 225 (2019) 119513.

[19] W. Yang, E. Fortunati, F. Bertoglio, J.S. Owczarek, G. Bruni, M. Kozanecki, J.M. Kenny, L. Torre, L. Visai, D. Puglia, Polyvinyl alcohol/chitosan hydrogels with enhanced antioxidant and antibacterial properties induced by lignin nanoparticles, Carbohydrate Polymers 181 (2018) 275–284.

[20] S.K. Sharma, A.P. Singh, In vitro antioxidant and free radical scavenging activity of Nardostachys jatamansi DC, J Acupunct Meridian Stud 5(3) (2012) 112–8.

[21] P.L. Thi, Y. Lee, D.L. Tran, T.T.H. Thi, J.I. Kang, K.M. Park, K.D. Park, In Situ Forming and Reactive Oxygen Species-Scavenging Gelatin Hydrogels for Enhancing Wound Healing Efficacy, Acta Biomater 103 (2020) 142–152.

[22] J. Li, C. Zhou, C. Luo, B. Qian, S. Liu, Y. Zeng, J. Hou, B. Deng, Y. Sun, J. Yang, Q. Yuan, A. Zhong, J. Wang, J. Sun, Z. Wang, N-acetyl cysteine-loaded graphene oxide-collagen hybrid membrane for scarless wound healing, Theranostics 9(20) (2019) 5839–5853.

[23] N. Ninan, A. Forget, V.P. Shastri, N.H. Voelcker, A. Blencowe, Antibacterial and Anti-Inflammatory pH-Responsive Tannic Acid-Carboxylated Agarose Composite Hydrogels for Wound Healing, ACS Appl Mater Interfaces 8(42) (2016) 28511–28521.

[24] H. Wu, F. Li, S. Wang, J. Lu, J. Li, Y. Du, X. Sun, X. Chen, J. Gao, D. Ling, Ceria Nanocrystals Decorated Mesoporous Silica Nanoparticle Based Ros-Scavenging Tissue Adhesive for Highly Efficient Regenerative Wound Healing, Biomaterials 151 (2018) 66–77.

[25] G. Hosgood, Wound healing. The role of platelet-derived growth factor and transforming growth factor beta, Vet Surg 22(6) (1993) 490–5.

[26] R. Noack, E. A. NEWSHOLME und C. START: REGULATION IN METABOLISM. 349 Seiten, 92 Abb., 57 Tab. John Wiley & Sons, London, New York, Sydney, Toronto 1973. Preis: 6.00 £, Food / Nahrung 19(7) (1975) 616–617.

[27] Z. Li, F. Wang, S. Roy, C.K. Sen, J. Guan, Injectable, highly flexible, and thermosensitive hydrogels capable of delivering superoxide dismutase, Biomacromolecules 10(12) (2009) 3306–16.

[28] S. Balaji, S.S. Vaikunth, S.A. Lang, A.Q. Sheikh, F.Y. Lim, T.M. Crombleholme, D.A. Narmoneva, Tissue-engineered provisional matrix as a novel approach to enhance diabetic wound healing, Wound Repair Regen 20(1) (2012) 15–27.

[29] H. Cho, S. Balaji, N.L. Hone, C.M. Moles, A.Q. Sheikh, T.M. Crombleholme, S.G. Keswani, D.A. Narmoneva, Diabetic wound healing in a MMP9-/-mouse model, Wound Repair Regen 24(5) (2016) 829–840.

[30] H. Cho, S. Balaji, A.Q. Sheikh, J.R. Hurley, Y.F. Tian, J.H. Collier, T.M. Crombleholme, D.A. Narmoneva, Regulation of endothelial cell activation and angiogenesis by injectable peptide nanofibers, Acta Biomater 8(1) (2012) 154–64.

[31] J.R. Hurley, H. Cho, A.Q. Sheikh, S. Balaji, S.G. Keswani, T.M. Crombleholme, D.A. Narmoneva, Nanofiber Microenvironment Effectively Restores Angiogenic Potential of Diabetic Endothelial Cells, Adv Wound Care (New Rochelle) 3(11) (2014) 717–728.

[32] H. Zhao, J. Huang, Y. Li, X. Lv, H. Zhou, H. Wang, Y. Xu, C. Wang, J. Wang, Z. Liu, Ros-Scavenging Hydrogel to Promote Healing of Bacteria Infected Diabetic Wounds, Biomaterials 258 (2020) 120286.

[33] B.S. Harrison, D. Eberli, S.J. Lee, A. Atala, J.J. Yoo, Oxygen producing biomaterials for tissue regeneration, Biomaterials 28(31) (2007) 4628–34.

[34] M.H. Lee, H.S. Jeon, S.H. Kim, J.H. Chung, D. Roppolo, H.J. Lee, H.J. Cho, Y. Tobimatsu, J. Ralph, O.K. Park, Lignin-Based Barrier Restricts Pathogens to the Infection Site and Confers Resistance in Plants, Embo j 38(23) (2019) e101948.

[35] Y. Hasegawa, E. Nakagawa, Y. Kadota, S. Kawaminami, Lignosulfonic acid promotes hypertrophy in 3T3-L1 cells without increasing lipid content and increases their 2-deoxyglucose uptake, Asian-Australas J Anim Sci 30(1) (2017) 111–118.

[36] M. Qiu, Q. Wang, Y. Chu, Z. Yuan, H. Song, Z. Chen, Z. Wu, Lignosulfonic Acid Exhibits Broadly Anti-HIV-1 Activity – Potential as a Microbicide Candidate for the Prevention of HIV-1 Sexual Transmission, PLOS ONE 7(4) (2012) e35906.

[37] S.C. Gordts, G. Férir, T. D’huys, M.I. Petrova, S. Lebeer, R. Snoeck, G. Andrei, D. Schols, The Low-Cost Compound Lignosulfonic Acid (LA) Exhibits Broad-Spectrum Anti-HIV and Anti-HSV Activity and Has Potential for Microbicidal Applications, PLOS ONE 10(7) (2015) e0131219.

[38] C. Huang, S. Tang, W. Zhang, Y. Tao, C. Lai, X. Li, Q. Yong, Unveiling the Structural Properties of Lignin–Carbohydrate Complexes in Bamboo Residues and Its Functionality as Antioxidants and Immunostimulants, ACS Sustainable Chemistry & Engineering 6(9) (2018) 12522–12531.

[39] Y. Zhang, M. Jiang, Y. Zhang, Q. Cao, X. Wang, Y. Han, G. Sun, Y. Li, J. Zhou, Novel Lignin-Chitosan-PVA Composite Hydrogel for Wound Dressing, Mater Sci Eng C Mater Biol Appl 104 (2019) 110002.

[40] D. Mahata, M. Jana, A. Jana, A. Mukherjee, N. Mondal, T. Saha, S. Sen, G.B. Nando, C.K. Mukhopadhyay, R. Chakraborty, S.M. Mandal, Lignin-graft-Polyoxazoline Conjugated Triazole a Novel Anti-Infective Ointment to Control Persistent Inflammation, Sci Rep 7 (2017) 46412.

[41] J. Xu, J.J. Xu, Q. Lin, L. Jiang, D. Zhang, Z. Li, B. Ma, C. Zhang, L. Li, D. Kai, H.D. Yu, X.J. Loh, Lignin-Incorporated Nanogel Serving as an Antioxidant Biomaterial for Wound Healing, ACS Appl Bio Mater 4(1) (2021) 3–13.

[42] T. Dizhbite, G. Telysheva, V. Jurkjane, U. Viesturs, Characterization of the radical scavenging activity of lignins--natural antioxidants, Bioresour Technol 95(3) (2004) 309–17.

[43] X. Pan, J.F. Kadla, K. Ehara, N. Gilkes, J.N. Saddler, Organosolv ethanol lignin from hybrid poplar as a radical scavenger: Relationship between lignin structure, extraction conditions, and antioxidant activity, J. Agric. Food Chem. 54 (2006) 5806−5813.

[44] S.B. Kedare, R.P. Singh, Genesis and development of DPPH method of antioxidant assay, J Food Sci Technol 48(4) (2011) 412–22.

[45] E.L.G. Samuel, M.T. Duong, B.R. Bitner, D.C. Marcano, J.M. Tour, T.A. Kent, Hydrophilic carbon clusters as therapeutic, high-capacity antioxidants, Trends Biotechnol 32(10) (2014) 501–505.

[46] S.A. Baba, S.A. Malik, Determination of total phenolic and flavonoid content, antimicrobial and antioxidant activity of a root extract of Arisaema jacquemontii Blume, Journal of Taibah University for Science 9(4) (2015) 449–454.

[47] D. Kai, W. Ren, L. Tian, P.L. Chee, Y. Liu, S. Ramakrishna, X.J. Loh, Engineering Poly(Lactide)– Lignin Nanofibers with Antioxidant Activity for Biomedical Application, ACS Sustainable Chem. Eng. 4(10) (2016) 5268.

[48] J. Wang, L. Tian, B. Luo, S. Ramakrishna, D. Kai, X.J. Loh, I.H. Yang, G.R. Deen, X. Mo, Engineering PCL/lignin nanofibers as an antioxidant scaffold for the growth of neuron and Schwann cell, Colloids Surf B Biointerfaces 169 (2018) 356–365.

[49] B. Das, P. Pal, P. Dadhich, J. Dutta, S. Dhara, In Vivo Cell Tracking, Reactive Oxygen Species Scavenging, and Antioxidative Gene Down Regulation by Long-Term Exposure of Biomass-Derived Carbon Dots, ACS Biomaterials Science & Engineering 5(1) (2019) 346–356.

[50] R. Liu, L. Dai, C. Xu, K. Wang, C. Zheng, C. Si, Lignin-Based Micro- and Nanomaterials and their Composites in Biomedical Applications, ChemSusChem 13(17) (2020) 4266–4283.

[51] J.A. Belgodere, D. Son, B. Jeon, J. Choe, A.C. Guidry, A.X. Bao, S.A. Zamin, U.M. Parikh, S. Balaji, M. Kim, J.P. Jung, Attenuating Fibrotic Markers of Patient-Derived Dermal Fibroblasts by Thiolated Lignin Composites, ACS Biomater Sci Eng 7(6) (2021) 2212–2218.

[52] S. Balaji, W.D. Short, B.W. Padon, J.A. Belgodere, S.E. Jimenez, N.T. Deoli, A.C. Guidry, J.C. Green, T.J. Prajapati, F. Farouk, A. Kaul, D. Son, O.S. Jung, C.E. Astete, M. Kim, J.P. Jung, Injectable Antioxidant and Oxygen-Releasing Lignin Composites to Promote Wound Healing, ACS Appl Mater Interfaces 15(15) (2023) 18639–18652.

[53] G. Carpentier, S. Berndt, S. Ferratge, W. Rasband, M. Cuendet, G. Uzan, P. Albanese, Angiogenesis Analyzer for ImageJ - A comparative morphometric analysis of “Endothelial Tube Formation Assay” and “Fibrin Bead Assay”, Sci Rep 10(1) (2020) 11568.

[54] P. Langfelder, S. Horvath, WGCNA: an R package for weighted correlation network analysis, BMC Bioinformatics 9 (2008) 559.

[55] P. Shannon, A. Markiel, O. Ozier, N.S. Baliga, J.T. Wang, D. Ramage, N. Amin, B. Schwikowski, T. Ideker, Cytoscape: a software environment for integrated models of biomolecular interaction networks, Genome Res 13(11) (2003) 2498–504.

[56] S.G. Keswani, A.B. Katz, F.Y. Lim, P. Zoltick, A. Radu, D. Alaee, M. Herlyn, T.M. Crombleholme, Adenoviral mediated gene transfer of PDGF-B enhances wound healing in type I and type II diabetic wounds, Wound Repair Regen 12(5) (2004) 497–504.

[57] M. Targosz-Korecka, G.D. Brzezinka, K.E. Malek, E. Stepień, M. Szymonski, Stiffness memory of EA.hy926 endothelial cells in response to chronic hyperglycemia, Cardiovasc Diabetol 12 (2013) 96.

[58] S. Costantino, F. Paneni, F. Cosentino, Hyperglycemia: a bad signature on the vascular system, Cardiovasc Diagn Ther 5(5) (2015) 403–6.

[59] G. Jin, Q. Wang, X. Pei, X. Li, X. Hu, E. Xu, M. Li, mRNAs expression profiles of high glucose-induced memory in human umbilical vein endothelial cells, Diabetes Metab Syndr Obes 12 (2019) 1249–1261.

[60] D.M. Nathan, P.A. Cleary, J.Y. Backlund, S.M. Genuth, J.M. Lachin, T.J. Orchard, P. Raskin, B. Zinman, Intensive diabetes treatment and cardiovascular disease in patients with type 1 diabetes, N Engl J Med 353(25) (2005) 2643–53.

[61] K. Thiem, S.T. Keating, M.G. Netea, N.P. Riksen, C.J. Tack, J. van Diepen, R. Stienstra, Hyperglycemic Memory of Innate Immune Cells Promotes In Vitro Proinflammatory Responses of Human Monocytes and Murine Macrophages, J Immunol 206(4) (2021) 807–813.

[62] F.R. González-Pacheco, J.J. Deudero, M.C. Castellanos, M.A. Castilla, M.V. Alvarez-Arroyo, S. Yagüe, C. Caramelo, Mechanisms of endothelial response to oxidative aggression: protective role of autologous VEGF and induction of VEGFR2 by H2O2, Am J Physiol Heart Circ Physiol 291(3) (2006) H1395–401.

[63] T. Kida, H. Oku, S. Osuka, T. Horie, T. Ikeda, Hyperglycemia-induced VEGF and ROS production in retinal cells is inhibited by the mTOR inhibitor, rapamycin, Sci Rep 11(1) (2021) 1885.

[64] K.E. Johnson, T.A. Wilgus, Vascular Endothelial Growth Factor and Angiogenesis in the Regulation of Cutaneous Wound Repair, Adv Wound Care (New Rochelle) 3(10) (2014) 647–661.

[65] A.J.M. Boulton, D.G. Armstrong, M.J. Hardman, M. Malone, J.M. Embil, C.E. Attinger, B.A. Lipsky, J. Aragón-Sánchez, H.K. Li, G. Schultz, R.S. Kirsner, Diagnosis and Management of Diabetic Foot Infections, American Diabetes Association, Arlington (VA), 2020.

[66] J. Folkman, E. Merler, C. Abernathy, G. Williams, Isolation of a tumor factor responsible for angiogenesis, J Exp Med 133(2) (1971) 275–88.

[67] D.W. Leung, G. Cachianes, W.J. Kuang, D.V. Goeddel, N. Ferrara, Vascular endothelial growth factor is a secreted angiogenic mitogen, Science 246(4935) (1989) 1306–9.

[68] A.P. Levy, N.S. Levy, S. Wegner, M.A. Goldberg, Transcriptional regulation of the rat vascular endothelial growth factor gene by hypoxia, J Biol Chem 270(22) (1995) 13333–40.

[69] J.A. Forsythe, B.H. Jiang, N.V. Iyer, F. Agani, S.W. Leung, R.D. Koos, G.L. Semenza, Activation of vascular endothelial growth factor gene transcription by hypoxia-inducible factor 1, Mol Cell Biol 16(9) (1996) 4604–13.

[70] N.M. Mazure, E.Y. Chen, K.R. Laderoute, A.J. Giaccia, Induction of vascular endothelial growth factor by hypoxia is modulated by a phosphatidylinositol 3-kinase/Akt signaling pathway in Ha-ras-transformed cells through a hypoxia inducible factor-1 transcriptional element, Blood 90(9) (1997) 3322–31.

[71] A.Y. Sheikh, J.J. Gibson, M.D. Rollins, H.W. Hopf, Z. Hussain, T.K. Hunt, Effect of hyperoxia on vascular endothelial growth factor levels in a wound model, Arch Surg 135(11) (2000) 1293–7.

