## Supplemental materials and figures for "Sustained ROS Scavenging and Pericellular Oxygenation by Lignin Composites Rescue HIF-1α and VEGF Levels to Improve Diabetic Wound Neovascularization and Healing"

#Corresponding authors

**Supplementary Methods**

RNA sequencing of MVEC

Extracted RNA underwent quantification using Qubit Fluorometer High sensitivity RNA assay (Thermo Fisher) and 500 ng of the extracted RNA was used for the library preparations and sequenced with the University of Houston Seq-N-Edit Core per standard protocols. RNAs were enriched and mRNA libraries were prepared with QIAseq® Stranded mRNA Library Kit. The size selection for libraries was performed using SPRIselect beads (Beckman Coulter) and purity of the libraries was analyzed using the High Sensitivity d1000 ScreenTape using Agilent Tapestation 4200. The prepared libraries were pooled and sequenced using Illumina NovaSeq 6000, generating ~30 million 2×150 bp paired end reads per sample. Raw sequencing data (Fastq) were loaded into CLC Genomics Workbench (Version 20.0.4 Qiagen) for data analysis. The Illumina sequencing adaptors were trimmed, and reads were mapped to the *mus musculus* ensemble v86 reference genome. Read alignment was represented as integer counts by using parameters of mismatch cost 2, insertion cost 3, deletion cost 3, length fraction 0.8, similarity fraction 0.8, max of 10 hits for a read. Integer read counts were normalized by Trimmed Means of M-values (TMM) algorithm.

Differential Gene Expression and Gene Ontology

To visualize differential gene expression (DGE), the FindAllMarkers function was employed with default settings to generate a volcano plot. Concurrently, gene ontology (GO) analysis was performed. Both analyses were conducted using RStudio (version 4.4.0) on the AnVIL platform. A basic volcano plot was created using the tidyverse package (version 1.3.0). From this plot, a threshold of |log_2_FC| > 0.5 and FDR p-value < 0.05 was established to identify differentially expressed genes. Genes were categorized as upregulated (log_2_FC > 0.5 and FDR p-value < 0.05), downregulated (log_2_FC < -0.5 and FDR p-value < 0.05), or not differentially expressed. The top 30 most significant genes were labeled on the volcano plot.

GO analysis was performed using the clusterProfiler package (version 4.12.1). A subset of genes with log_2_FC values greater than 0.5 was extracted from the FindAllMarkers data for this analysis. Additional dot plots were generated to visualize the relationship between gene ratio and the number of genes associated with each GO term.

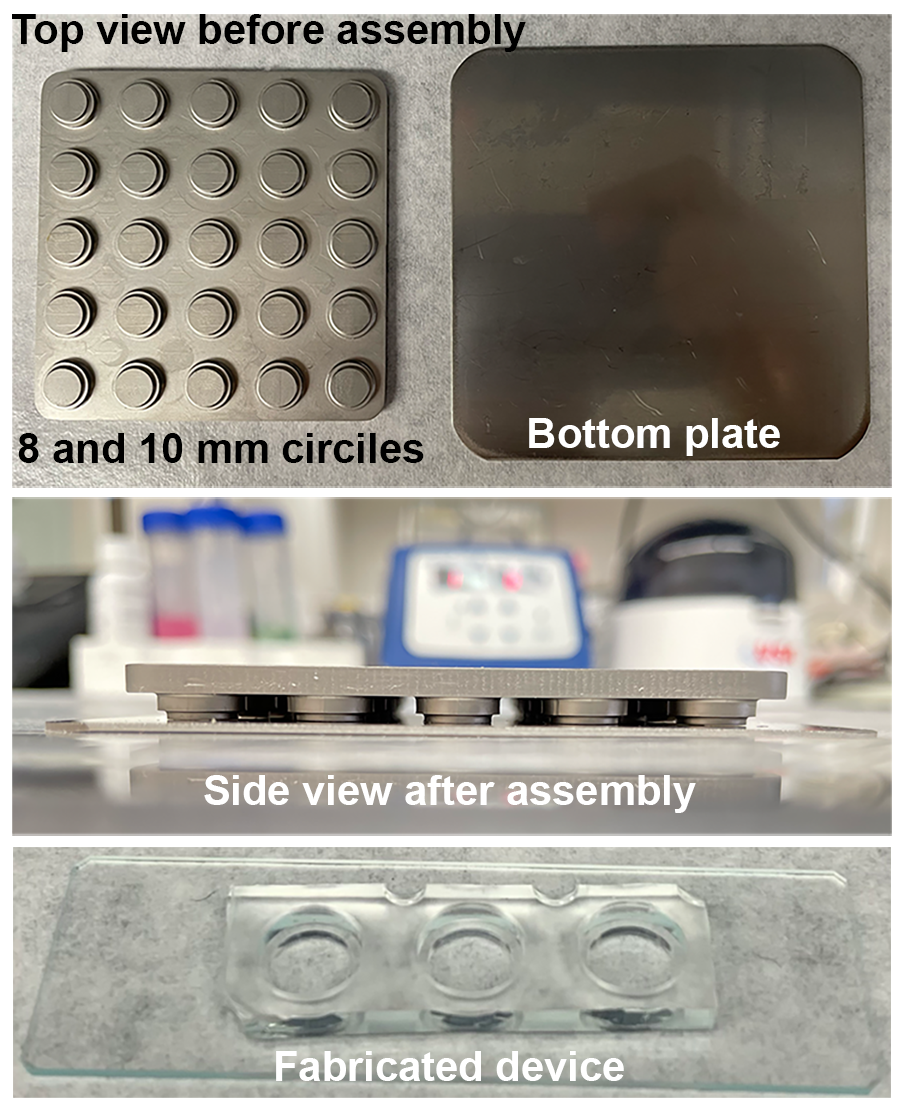
**Supplementary Figures**

**Figure S1.** Precision metal plate (positive mold) for PDMS (Sylcap 284-F) wells. The metal bottom plate is used to ensure a flat surface for enhanced plasma bonding of PDMS to a glass surface. Lignin composite wells (8 mm diameter; 1 mm height) and medium wells (10 mm diameter; 2 mm height) are fabricated with the metal mold. PDMS is cured with the positive mold in a rectangular petri dish.

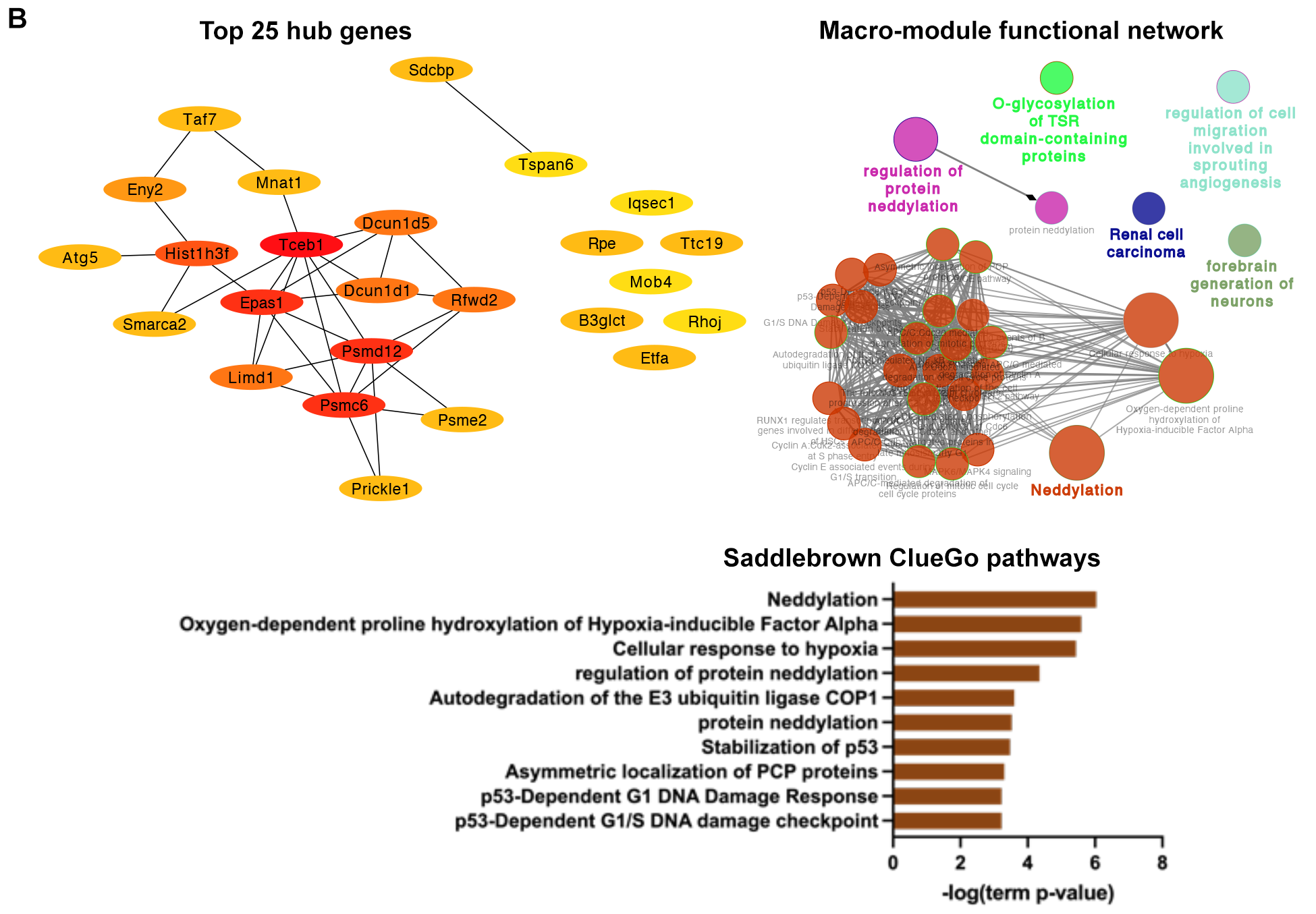

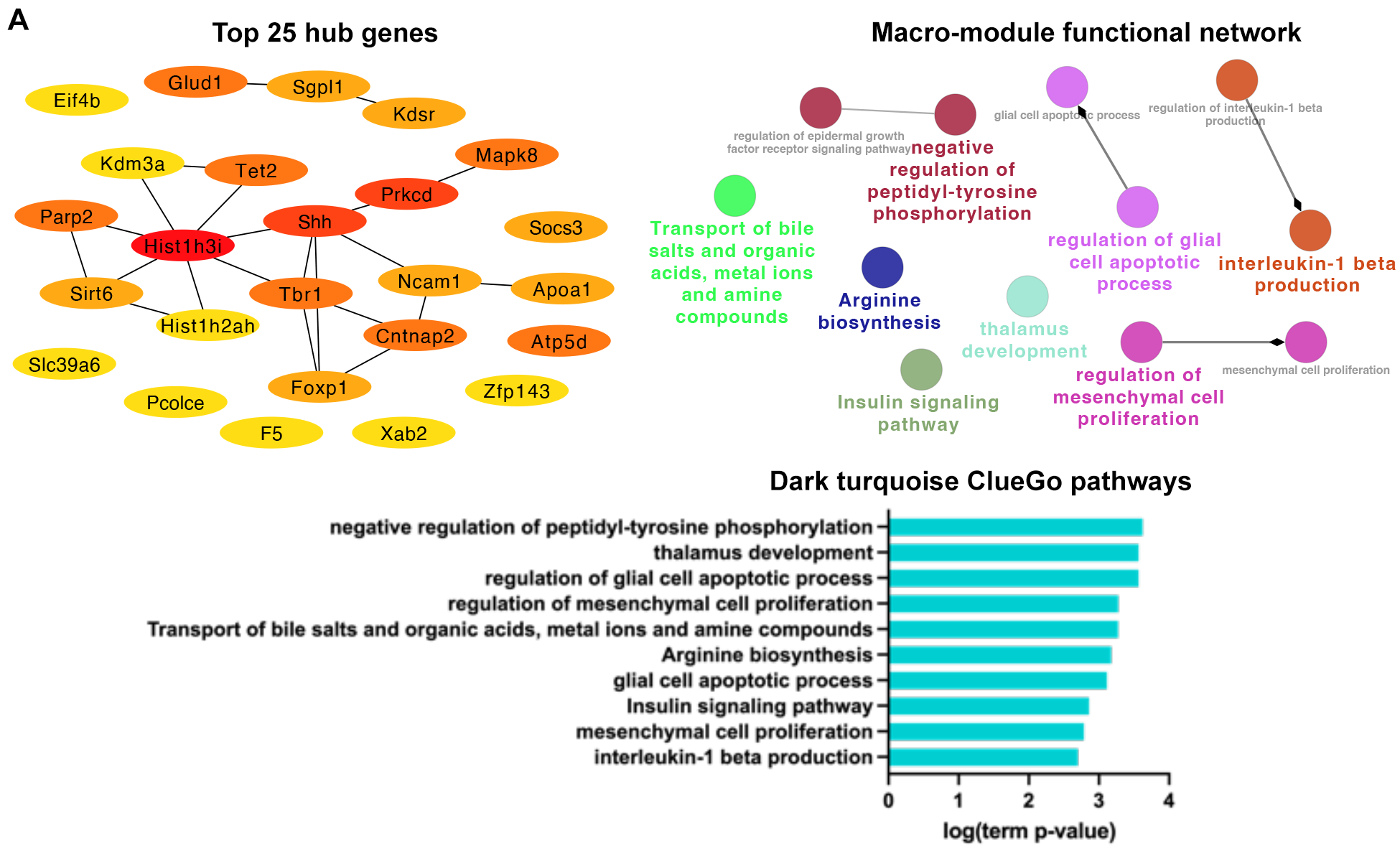

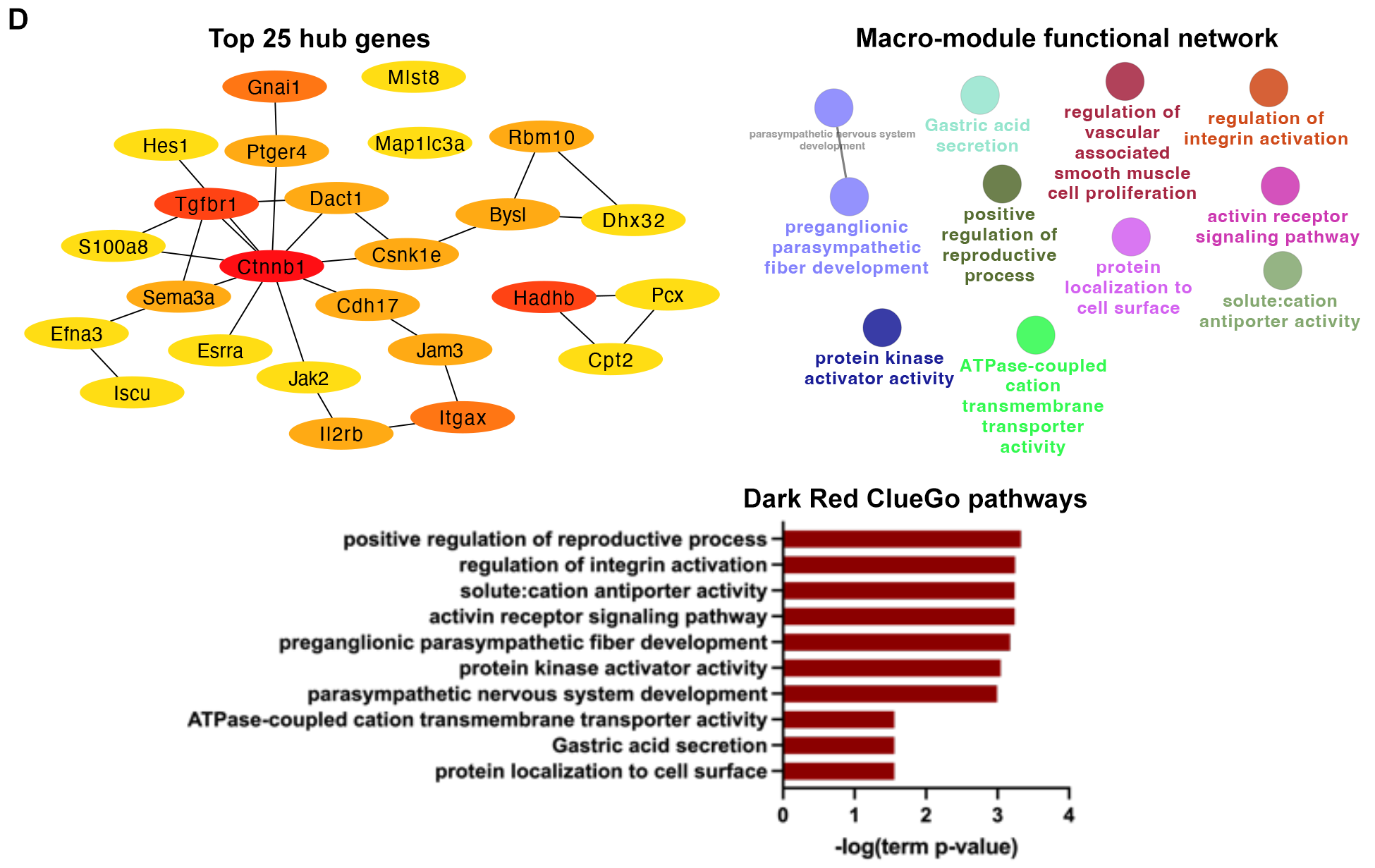

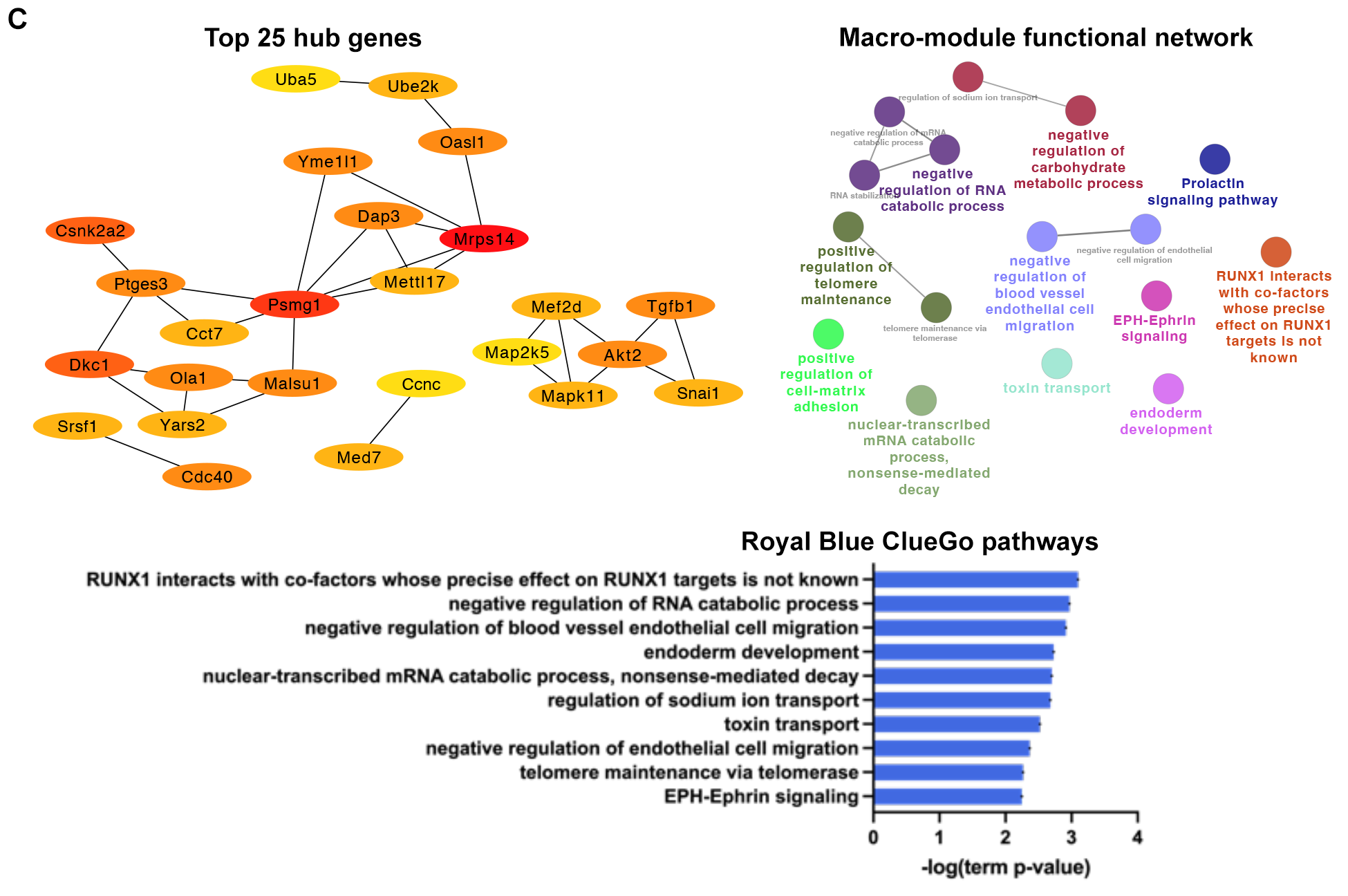

**Figure S2.** Top 25 hub genes identified in each additional module discovered through WGCNA. Macro-module functional relationships and ClueGO pathway enrichment analyses are also presented. In addition to the main turquoise module, dark turquoise (191 genes, A), saddle brown (135 genes, B), royal brown (127 genes, C), dark red (201 genes, D) and black (1403 genes, E) modules are selected from **Figure 3B** module-sample relationship.

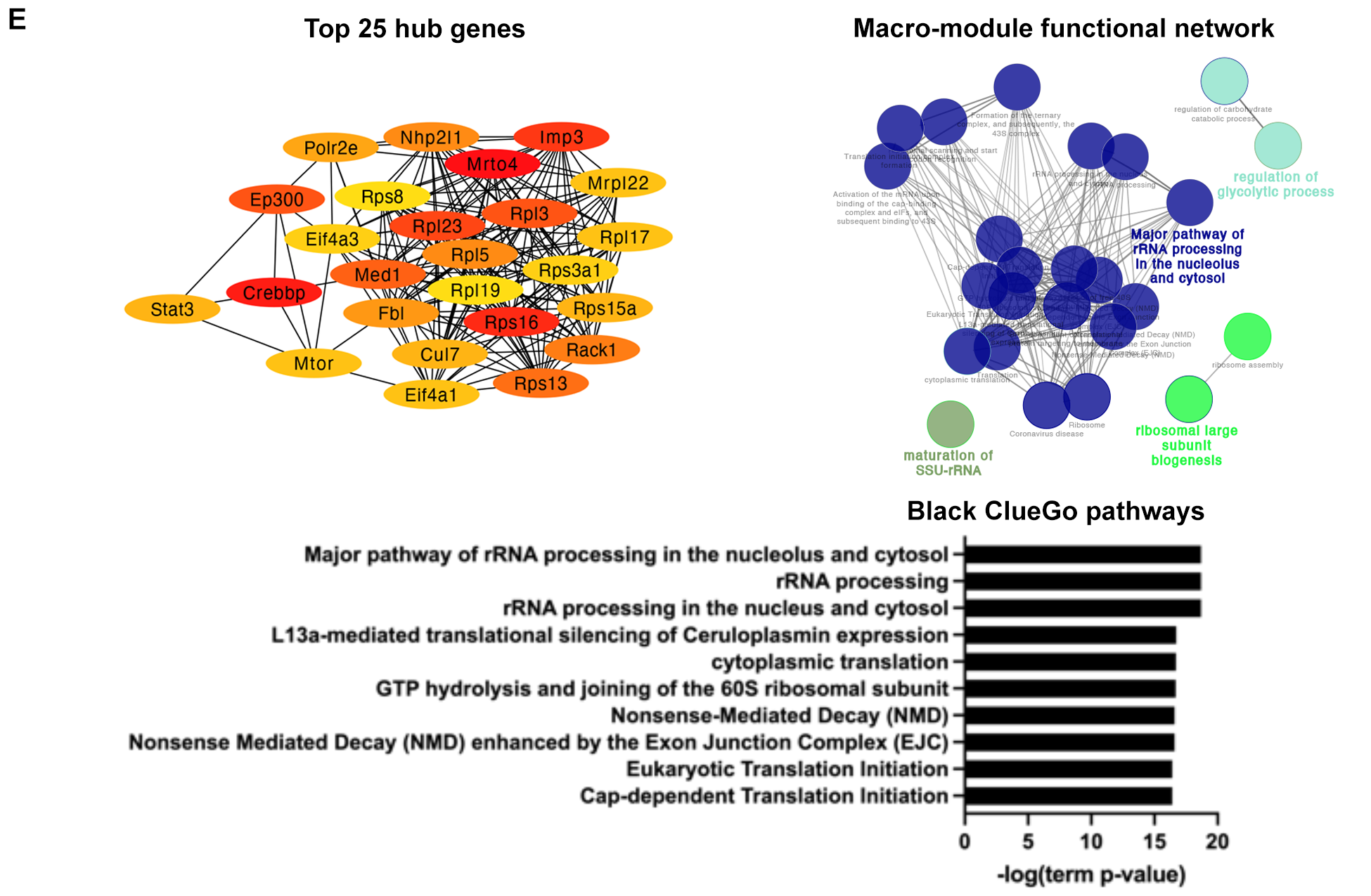

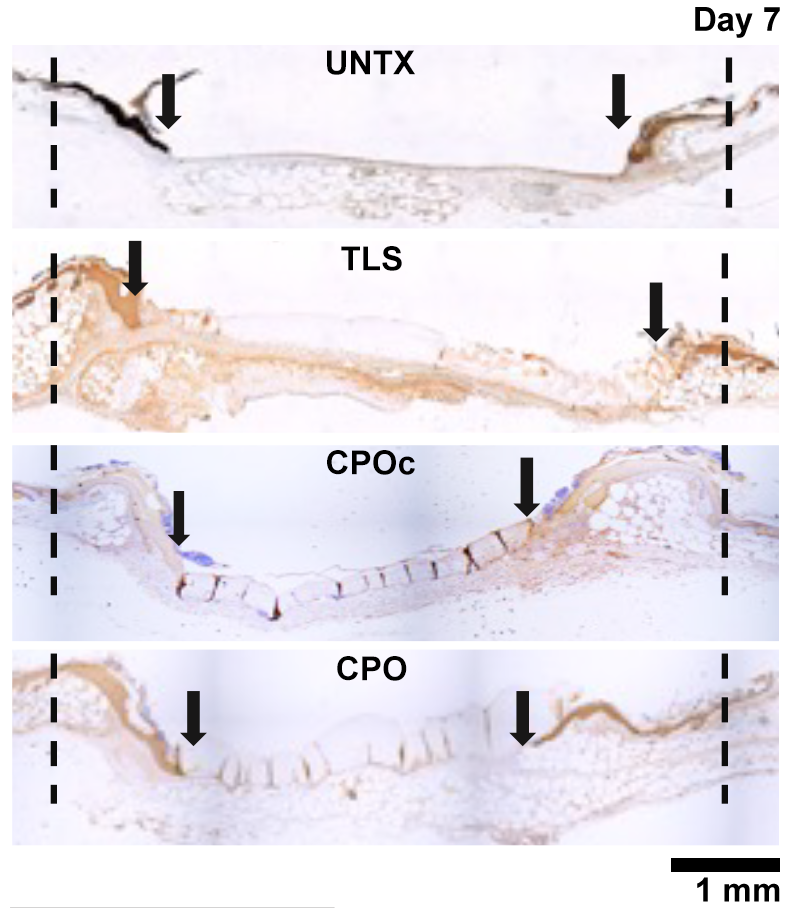

**Figure S3.** (A) Representative cK14 (cytokeratin 14) staining of post-wounding day 7 wounds, Scale bar = 1 mm.

**Figure S4.** Healing progression of db/db mouse wounds treated with lignin composite TLS, CPO or CPOc immediately post-wounding at 14 days post-wounding as compared to UNTX wounds. (A) Gross images taken at time of tissue collection at day 14 post-wounding, with representative H&E staining of the wound sections. Scale bar = 500 µm. (B) Staining of day 14 wound sections with antibodies against CD31. Scale bar = 50 µm. (C) CD31 staining of wound sections showed an increase in visible lumens in the CPO lignin composite treated group at day 14. Mean±SD, n=3 wounds per treatment.

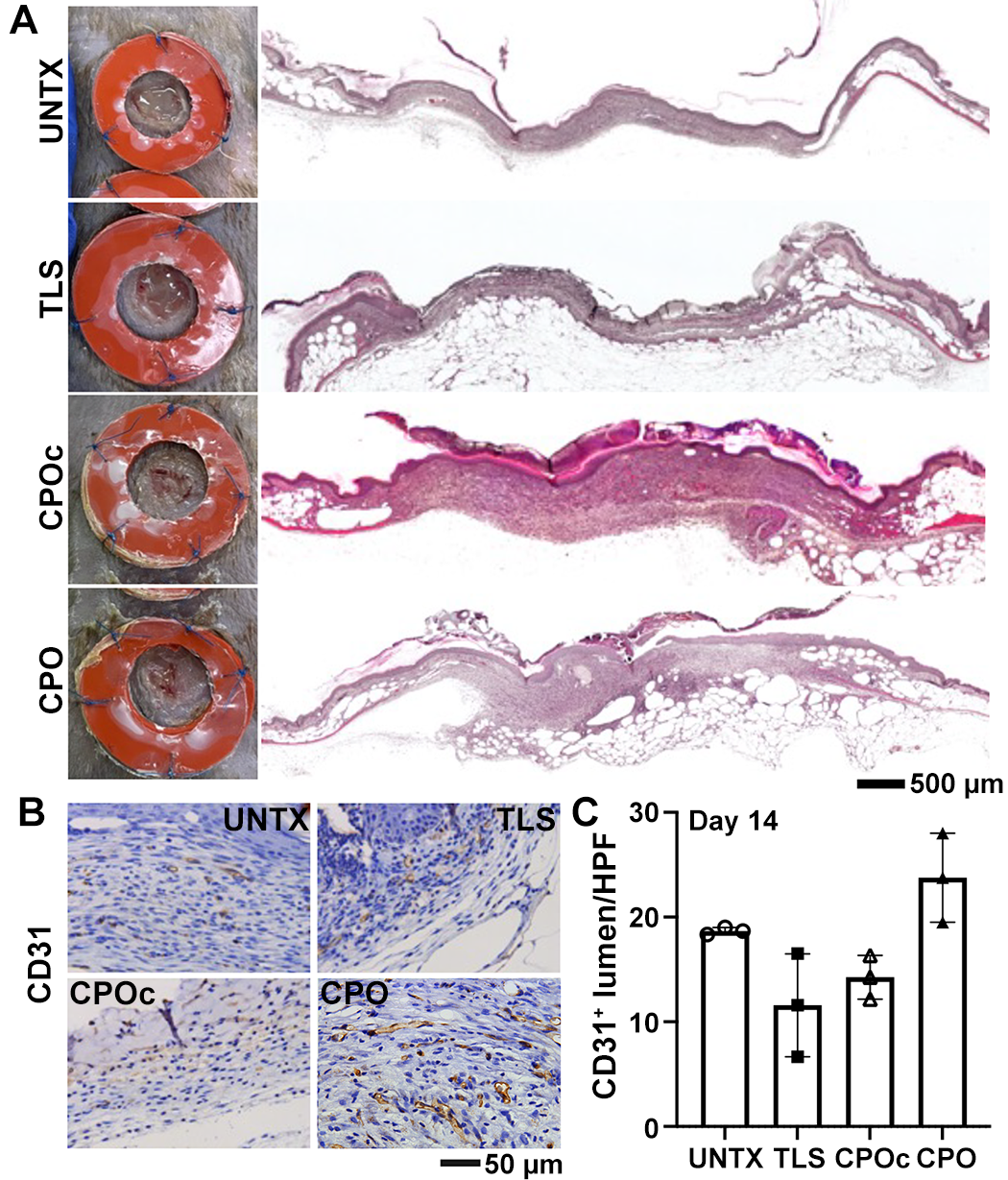

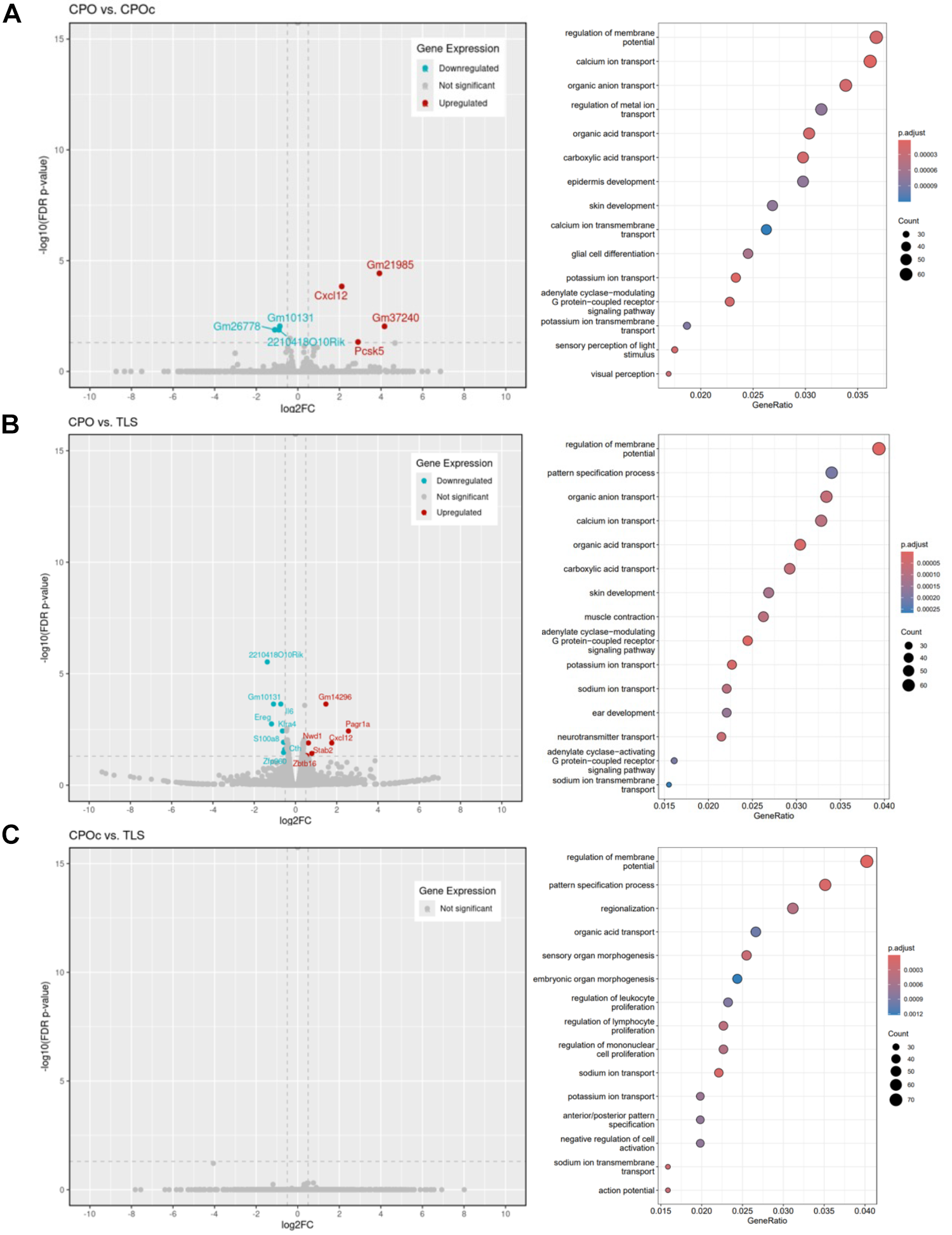

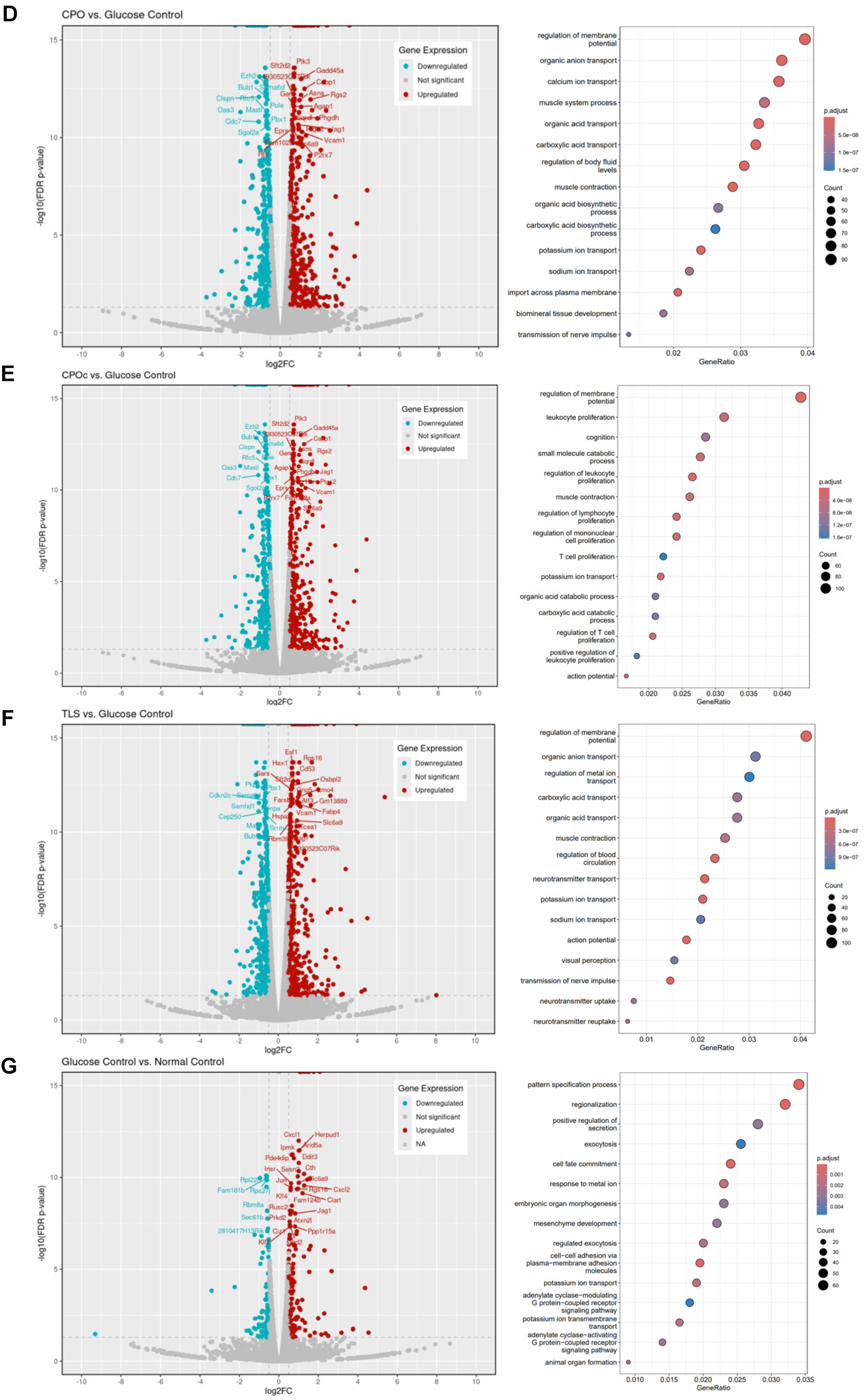
**Figure S5.** DGE and GO of samples: CPO vs. CPOc (A), CPO vs TLS (B), CPOc vs TLS (C), CPO vs. GelMA (glucose) (D), CPOc vs GelMA (glucose) (E), TLS vs GelMA (glucose) (F) and GelMA (glucose) vs GelMA (normal) (G).
